# The Barley HvSTP13GR mutant triggers resistance against biotrophic fungi

**DOI:** 10.1101/2021.09.16.460598

**Authors:** Ines Caroline Skoppek, Wilko Punt, Marleen Heinrichs, Frank Ordon, Gwendolin Wehner, Jens Boch, Jana Streubel

## Abstract

High-yielding and stress resistant crops are essential to ensure future food supply. Barley is an important crop to feed livestock and to produce malt, but the annual yield is threatened by pathogen infections. Pathogens can trigger an altered sugar partitioning in the host plant, that possibly leads to an advantage for the pathogen. Hampering these processes represents a promising strategy to potentially increase resistance. We analyzed the response of the barley monosaccharide transporter *HvSTP13* towards biotic stress and its potential use for plant protection. The expression of *HvSTP13* increased upon bacterial and fungal PAMP application, suggesting a PAMP-triggered signaling that converged on the transcriptional induction of the gene. Promoter studies indicate a region that is likely targeted by transcription factors downstream of PAMP-triggered immunity pathways. We confirmed that the non-functional HvSTP13GR variant confers resistance against an economically relevant biotrophic rust fungus, in barley. In addition, we established targeted CRISPR/Cas9 cytosine base editing in barley protoplasts to generate alternative *HvSTP13* mutants and characterized the sugar transport activity and subcellular localization of the proteins. These mutants represent promising variants for future resistance analysis. Our experimental setup provides basal prerequisites to further decode the role of *HvSTP13* in response to biological stress. Moreover, in line with other studies, our experiments indicate that the alteration of sugar partitioning pathways, in a host pathogen interaction, is a promising approach to achieve broad and durable resistance in plants.

## 1 Introduction

Barley is not only one of the major crops to feed livestock and an essential ingredient of beer, it has also gained a growing importance in food production. However, the infection with fungal pathogens is one of the most devastating threats to annual yields by causing severe damages of various organs (Gaj *et al*., 2016; Liu *et al*., 2011; Park *et al*., 2015; Wolfe and McDermott, 2003). To ensure a sustainable and high-yield crop production, an ongoing improvement towards resistant varieties is therefore the most important challenge for research and plant breeding.

An ecologically sustainable way of crop protection is the identification of and breeding with naturally occurring resistance genes, that originate in related wild varieties or landraces (Dinh *et al*., 2020). However, many resistances are mainly effective against certain races of one fungal species or have already been overcome (Clifford, 1985; König *et al*., 2012; Niks *et al*., 2000). The identification of new resistance mechanisms is therefore a major future challenge. One promising approach is to eliminate fundamental requirements for a successful pathogen infection. The availability of sugars to cover their energy demand is a major need for plant pathogens (Bezrutczyk *et al*., 2018). Thus, uncovering the routes of nutrient utilization by fungal pathogens and developing methods that block feeding, could represent a highly promising strategy to engineer a durable and broad-spectrum resistance.

Sugars are the basal energy source for living organisms and their acquisition and partitioning are key steps to conduct physiological processes. In plants, sugars are produced in photosynthetically active source tissues and distributed to supply energy to consuming sink tissues. Specialized sugar transporters are crucial players in these delivery pathways and belong to three major groups: the SWEETs, SUTs and STPs. Members of the SWEET-family typically consist of seven transmembrane (TM) domains and transport a variety of substrates. Sucrose H+ co-transporters (SUTs) and sugar transport proteins (STPs) belong to the major facilitator superfamily (MFS) with typically 12 TM domains (Büttner, 2010; Julius *et al*., 2017). Sucrose exporting SWEETs and SUTs control the key step of phloem loading and long distance transport. STPs, on the other hand, transport monosaccharides, like glucose or fructose, over short distances, thereby providing ready-to-use sugars to feed energy consuming cells and cell compartments (Büttner, 2010; Eom *et al*., 2015; Julius *et al*., 2017).

As the low sugar content in the plant apoplast might constrain their growth, pathogens utilize the sugar reservoirs of their host to convert the infection area into an artificial sink (Pommerrenig *et al*., 2020). Especially SWEETs are well known targets that are exploited by diverse fungal and bacterial pathogens, e.g. *Botrytis cinerea, Blumeria graminis, Ustilago maydis, Pseudomonas syringae* and various *Xanthomonas* species (Breia *et al*., 2020; Chen *et al*., 2010; Doidy *et al*., 2012; Sosso *et al*., 2019; Timilsina *et al*., 2020). In response, plants can activate the expression, or enhance the transport activity of SUTs or STPs to counter the pathogen-induced leakage of sugars (Bezrutczyk *et al*., 2018; Pommerrenig *et al*., 2020).

One of the most prominent STPs involved in plant-pathogen interactions is STP13, but its role for the outcome of the respective interaction seems a double-edged sword. On the one hand, an increased STP13 activity supports basal resistance, as shown for the interaction of the necrotrophic fungus *Botrytis cinerea* with *A. thaliana*. Hence, it was hypothesized that the increased *AtSTP13* expression might remove glucose from the infection site to slow down the infection process (Lemonnier *et al*., 2014). Notably, *AtSTP13* is also involved in interactions with bacterial pathogens. Upon detection of flg22 by the plant receptor FLS2, *AtSTP13* is expressed and phosphorylated by BAK1 (Yamada *et al*., 2016). This modification enhances the sugar uptake of the transporter, possibly causing it to compete with the pathogen for sugar availability in the apoplast.

In contrast, an overexpression of the glucose transporter TaSTP13 supports the virulence of biotrophic fungi in wheat and Arabidopsis whereas silencing of *TaSTP13* in wheat increased resistance (Huai *et al*., 2020). Because biotrophic fungi feed from living cells, an increased STP13 activity might enhance the glucose availability in the plant cytoplasm and thereby feed the pathogens via their haustoria. In line with this assumption is the observation that the glucose transport-deficient TaLr67res (G144 to R144 exchange, hereafter named TaSTP13GR) variant from wheat increases the resistance of wheat and *TaSTP13GR*-transgenic barley against the biotrophic fungi *Puccinia hordei* and *Blumeria graminis* (Milne *et al*., 2019; Moore *et al*., 2015). Moreover, just recently, it was shown that a non-functional MtSTP13 G to R variant also enhances the resistance of *Medicago truncatula* to the biotrophic powdery mildew fungus *Erysiphe pisi* (Gupta *et al*., 2021). This suggests, that an impeded glucose import from the plant apoplast to the cytoplasm might starve the fungal haustoria, thus preventing progression of a severe infection.

The barley ortholog *HvSTP13* harboring the G144 to R144 mutation (hereafter named HvSTP13GR) also lacks glucose transport activity but it was not analyzed whether this mutant can confer resistance against biotrophic pathogens, like it was observed for TaSTP13GR (Milne *et al*., 2019). If that is the case, it would open the way for barley plants that are resistant against a broad spectrum of biotrophic fungi.

In this study we analyzed the expression pattern of the barley hexose transporter *HvSTP13* during various biotic stresses. We observed that *HvSTP13* is induced in response to several fungal and bacterial pathogens as a result of the recognition of certain PAMPs. By using GUS reporter studies we dissected the *HvSTP13* promoter and identified a region that is likely responsible for the induced expression. With transgenic plants we analyzed the effect of the HvSTP13GR mutant and found that it confers resistance against an economically relevant biotrophic rust fungus. In addition, a CRISPR/Cas9 cytosine base editing approach, in barley protoplasts, identified alternative *HvSTP13* mutants as new candidates for future resistance analysis. In summary our study contributes new insights into the molecular trigger for *HvSTP13* expression and its potential to confer broad spectrum resistance against biotrophic pathogens.

## 2 Results

### 2.1 The expression of *HvSTP13* is induced by fungal and bacterial PAMPs

In barley, *HvSTP13* is induced upon treatment with the leaf rust fungus *Puccinia hordei* and the powdery mildew fungus *Blumeria graminis* pv. *hordei* (Milne *et al*., 2019). To further dissect the *HvSTP13* expression pattern under diverse biotic pressures, we infected barley seedlings with the biotrophic fungi *Puccinia hordei* (*Ph*), *Puccinia striiformis* f. sp. *hordei* (*Psh*) and *Blumeria graminis* pv. *hordei* (*Bgh*), or with the necrotrophic fungus *Pyrenophora teres* pv. *teres (Ptt)*. Leaf samples were harvested at 24, 48 and 72 hours post inoculation (hpi) and total RNA was extracted for qRT-PCR. The infection was confirmed by analyzing the expression of a plant chitinase (*HvPR3*) that is known to respond to fungal infection (Figure S1; (Ali *et al*., 2018)). In contrast to mock inoculated plants, all four fungal pathogens elicited an increase in *HvSTP13* expression between 24 and 72 hpi (Figure 1a). To analyze, whether *HvSTP13* is also induced upon bacterial treatment, we inoculated the host pathogen *Xanthomonas translucens* pv. *hordei* UPB820 (*Xth*), as well as the non-host pathogen *Pseudomonas syringae* pv. *tomato* DC3000 (*Pto*) into barley and found an increase of *HvSTP13* expression for both bacterial pathogens (Figure 1b). A detailed time curve, performed with UPB820, indicates that *HvSTP13* induction starts between 1 and 4 hpi (Figure S2a). Next, we analyzed the expression of *HvSTP13* in barley after inoculation with the PAMPs chitin, flg22, β-1,3-glucan and laminarin. An increase of *HvSTP13* transcript was detected with flg22 and chitin treatment not later than 4 hpi (Figure 1c). In contrast, laminarin and β-1,3-glucan did not activate the expression of *HvSTP13* (Figure S2b). These results confirm, not only, that *HvSTP13* is induced as a consequence of fungal and bacterial PAMP-signaling, they also suggest a potential role in PTI-related processes.

**Figure 1.**
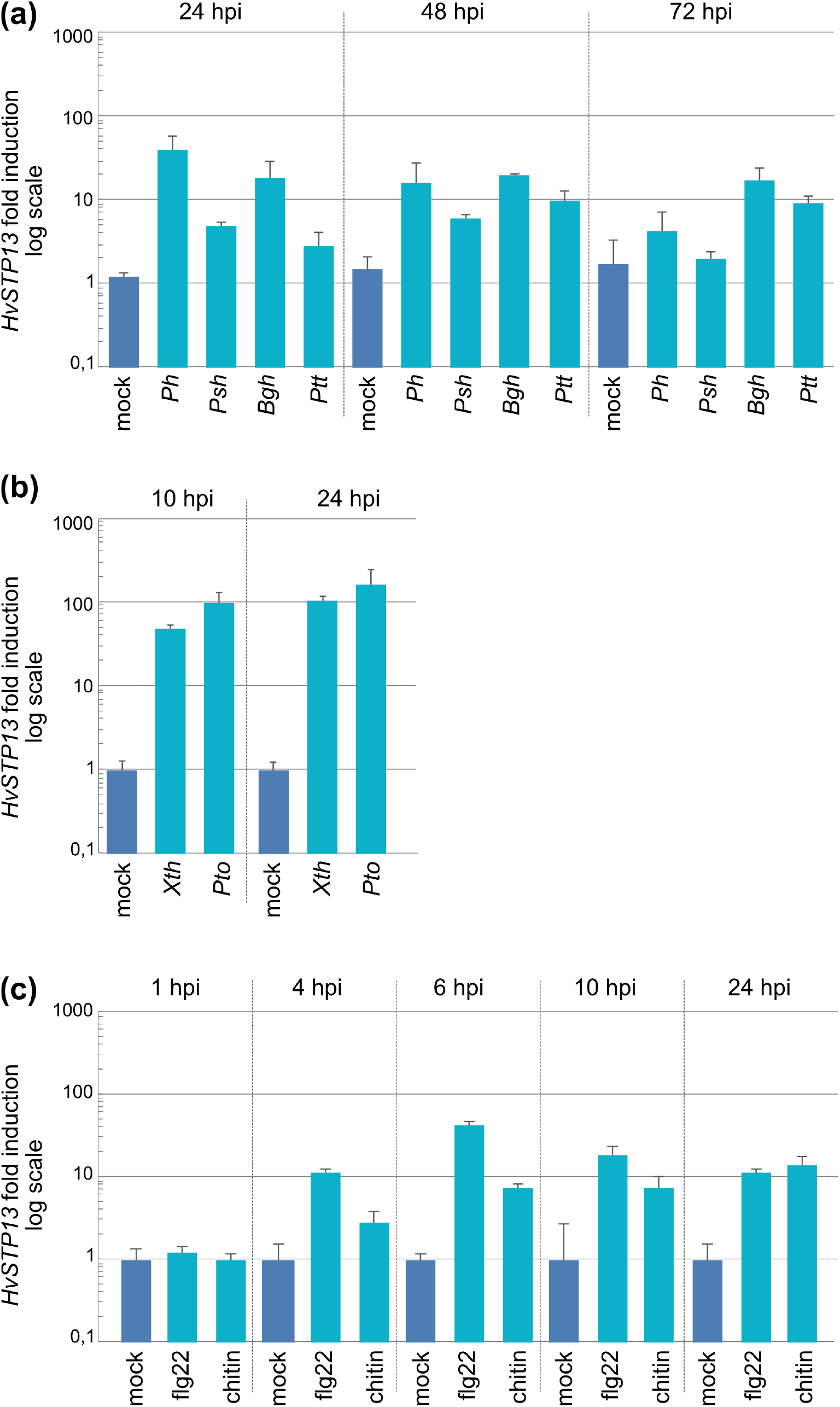
Transcript level of *HvSTP13* after treatment with pathogens or PAMPs, respectively. Barley plants were inoculated with **(a)** *Puccinia hordei (Ph*), *Puccinia striiformis* f. sp. *hordei* (*Psh*), *Blumeria graminis* pv. *hordei* (*Bgh*) and *Pyrenophora teres* pv. *teres* (*Ptt*) **(b)** *Xanthomonas translucens* pv. *hordei* strain UPB820 (*Xth*) and *Pseudomonas syringae* pv. *tomato* strain DC3000 (*Pto*) or **(c)** with the PAMPs flg22 and chitin. **(a) – (c)** Each bar represents three biological replicates. Mock = inoculation medium; hpi: hours post inoculation. Error bars represent the standard deviation.

### 2.2 The promoter of *HvSTP13* harbors PAMP-responsive elements

In order to identify elements that might be required for the pathogen-mediated transcriptional activation, we used PLACE and PlantCARE to predict cis-regulatory elements (CREs) in an approximately 2 kb region of the *HvSTP13* promoter (Higo *et al*., 1998; Lescot *et al*., 2002). In accordance to the response of the *STP13* expression to multifaceted biotic and abiotic factors, the *in silico* analysis predicted a variety of different CREs within the *HvSTP13* promoter region (Table S1 & S2). Abscisic acid (ABA) has been shown to activate the expression of *STP13* in Arabidopsis, barley, wheat, grapevine and *Medicago truncatula* (Gupta *et al*., 2021; Hayes *et al*., 2010; Huai *et al*., 2020; Milne *et al*., 2019; Yamada *et al*., 2011). Consistent with this, the prediction revealed the presence of several ABA-responsive elements (ABREs) in the *HvSTP13* promoter. Additionally, the *in silico* prediction identified several promising binding sites for WRKY transcription factors (W-box) and one sugar responsive element (SURE) – two CRE types that are involved in immunity and sugar signaling pathways (Table S1 & S2; Figure S3a (Chen *et al*., 2019; Sun *et al*., 2003)).

To analyze which promoter regions are required for the PAMP-triggered induction of *HvSTP13*, we amplified four truncated *HvSTP13* promoter variants that lack certain CREs. We labeled them according to the number of nucleotides upstream of the transcription start site (TSS). This results in promoter fragment p-887, p-445, p-222 and p-93. All used promoter fragments additionally included the 301 bp of the 5’ UTR (Figure 2a & S3a). The promoter variants were fused to a promoterless GUS reporter gene and transformed into barley embryos to generate transgenic GUS reporter lines. For each construct, three independent T1 lines, with six plants each, were analyzed. The presence of the transgene was verified by PCR. Negative siblings (indicated with -), as well as wildtype Golden Promise plants, were used as negative controls. Leaf discs were either infiltrated with flg22, chitin or MgCl_2_. The resulting GUS activity was analyzed after 24 hours. The two fragments *p-887* and *p-445* displayed an increase of GUS activity after treatment with flg22 and chitin, indicating an induction after PAMP signaling. For the two shorter fragments, *p-222* and *p-93*, no difference in GUS activity was observed between transgenic and non-transgenic siblings after flg22 or chitin treatment (Figure 2b). This indicates that the region between -445 and -222 bp upstream of the TSS possibly contains important cis-regulatory elements, which could be bound by regulatory transcription factors, and thus allow for PAMP-mediated induction.

**Figure 2.**
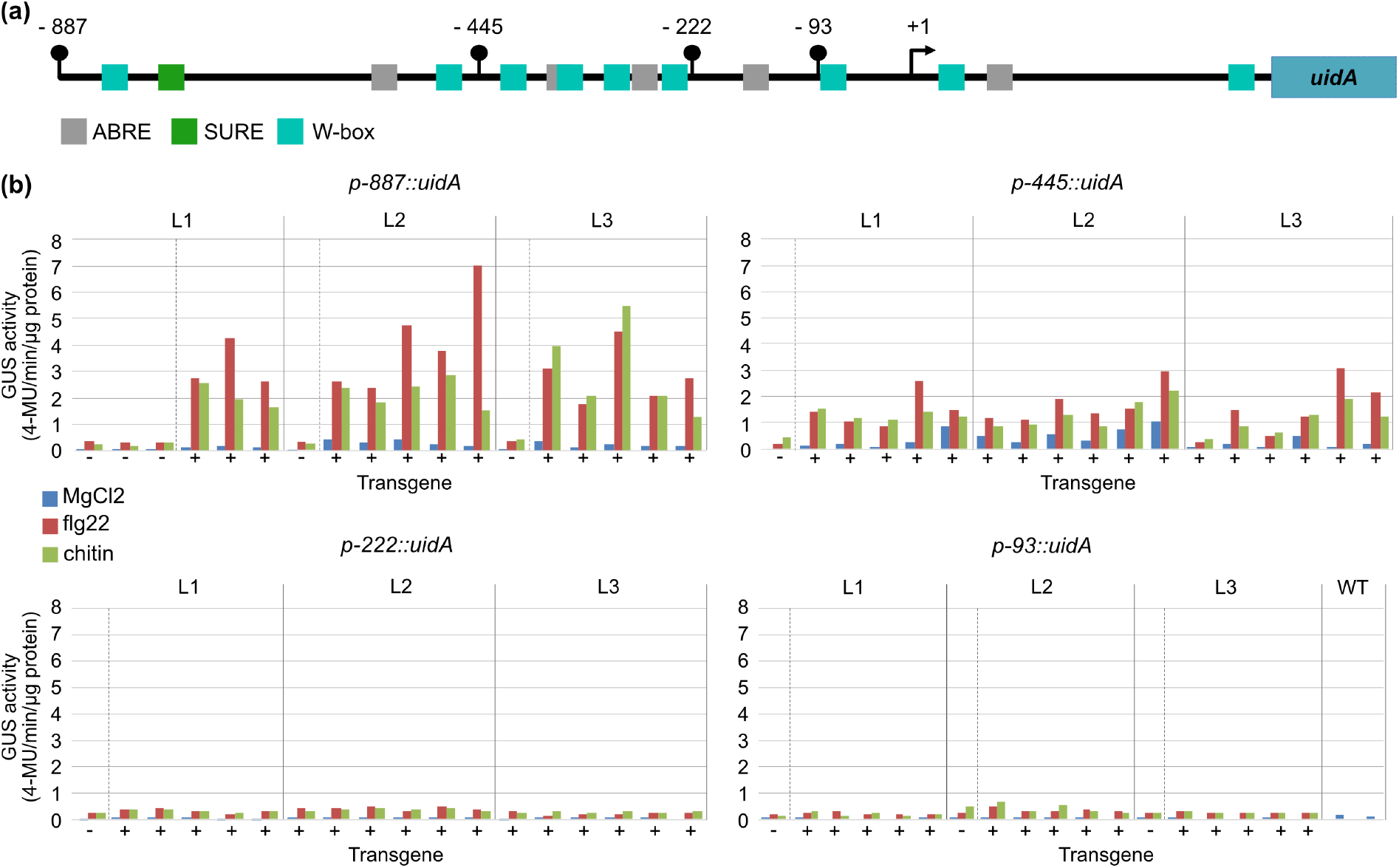
Analysis of the PAMP-triggered activation of truncated *HvSTP13* promoter fragments. **(a)** Schematic overview of the truncated *pHvSTP13* fragments. The black arrow marks the transcription start site (+1) as annotated in Phytozome. The length of each truncated promoter fragment is marked by a black dot. Predicted ABREs (grey), SURE (green) or W-boxes (turquoise) are indicated. The truncated promoter fragments were fused to the *uidA* reporter gene. **(b)** For each reporter construct six individual plants of three T1 lines (L1, L2, L3) were used. For each plant the presence of the transgene was analyzed by PCR (-no transgene; + transgenic). Leaf discs of each plant were vacuum infiltrated with PAMPs or MgCl_>2_>. GUS activity was measured 24 hpi. Each bar represents the mean of the technical replicate.

### 2.3 New HvSTP13 mutants can be achieved by cytosine base editing in barley protoplasts

STP13GR variants are known to confer fungal resistance when introduced as a transgene. It would be highly valuable to introduce such a mutation into the genomic copy of barley *HvSTP13* by using new breeding techniques. Unfortunately, CRISPR Cas9 based cytosine or adenine base editors are not suitable to exactly convert the triplet for glycine at position 144 to arginine. Therefore, we scanned the HvSTP13 sequence to identify amino acids that might similarly impact the sugar transport activity and which can be targeted by CRISPR/Cas9 base editors. One promising target is amino acid D41. This amino acid is equivalent to D42 in the sugar transporter ortholog AtSTP10. In AtSTP10, D42 forms a proton donor-acceptor pair with R142 to translocate an H+ proton (Paulsen *et al*., 2019). The transport of the proton is coupled with the glucose transport and a mutation in AtSTP10 D42 results in a loss of glucose transport activity in AtSTP10 (Paulsen *et al*., 2019). Possibly, a mutation of D41 in HvSTP13 might also result in a loss of H+ translocation and thus in a loss of glucose transport activity.

To achieve D41 mutants by base editing, we constructed a cytosine base editor with a wheat codon optimized nCas9 and a sgRNA targeting the nucleotide sequence surrounding the triplet for D41 (Figure S4a). The sgRNA expression was controlled by the *TaU6* promoter. It was either used as a single unit or as a consecutive repetition of four expression units, to increase sgRNA abundance (Figure S4a). We analyzed the editing results by amplicon sequencing following two independent transformations of barley protoplasts. In both repetitions, we detected sequence variants that converted the three amino acid stretch D41, V42 and G43 to N41, I42 and S43 or N43 (NIS or NIN; Figure S4b,c). Additionally, we observed a conversion of two amino acids to N41 and I42 (NI; Figure S4b,c) or the single D41 to N41 mutant (N; Figure S4b,c). This approach confirmed that it will, likely, be possible to generate D41 mutations using base editing in barley cells and likely also whole plants.

### 2.4 Mutant variants of *HvSTP13* are localized in the plasma membrane but lack glucose transport activity

To analyze the glucose transport activity of the D41 HvSTP13 variants we constructed the *HvSTP13NIS* triple and the *HvSTP13N41* single mutant (Figure 3 & 4). As reference we included the *HvSTP13* wildtype and the *HvSTP13GR* mutant (Milne *et al*., 2019). All *HvSTP13* variants were cloned into yeast expression vectors and transformed into the yeast strain EBY.VW4000 (Wieczorke *et al*., 1999). This strain lacks 18 hexose and 3 additional transporter genes and does not grow on glucose, fructose, galactose or mannose as carbon source. The growth of EBY.VW4000 on maltose is not affected. Thus, EBY.VW4000 is frequently used to analyze the hexose transport function of heterologous proteins. As positive control we included the yeast hexose transporter *Hxt1* that is known to restore growth of EBY.VW4000 on glucose containing medium. *eGFP* served as negative control (Wieczorke *et al*., 1999). The resulting yeast strains were spotted on selective medium containing either glucose or maltose as carbon source. On maltose containing media, all strains showed comparable growth, thus providing a growth control (Figure 3). However, on glucose containing media, only the heterologous expression of the wildtype *HvSTP13* transporter and the positive control *Hxt1* conferred growth. In contrast, neither *HvSTP13GR* nor the new *HvSTP13* mutants or the negative control *eGFP* conferred growth on glucose containing media. This demonstrates that only *HvSTP13* complemented the glucose uptake deficiency of the yeast strain EBY.VW4000, whereas the new HvSTP13 D41 mutants are just as incapable to transport glucose as HvSTP13GR.

**Figure 3.**
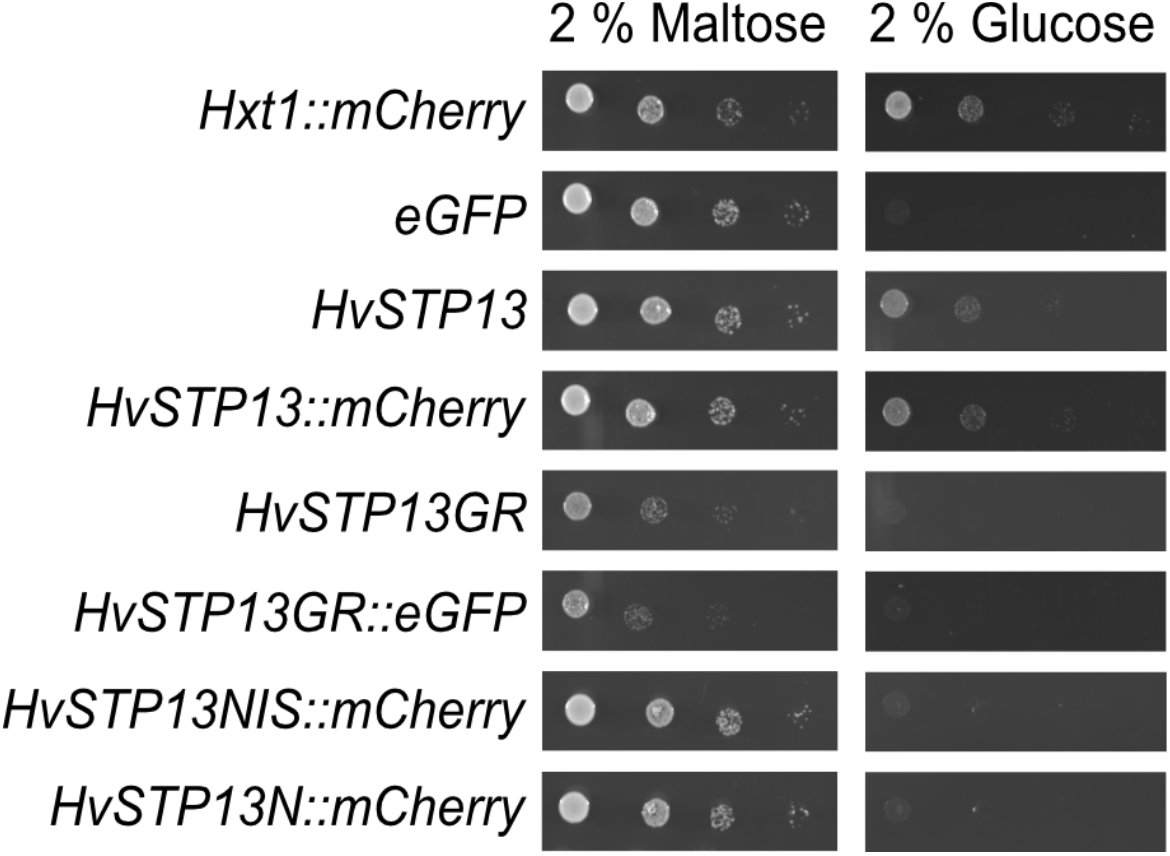
Transport analysis of *HvSTP13* variants in the hexose transport-deficient strain EBY.VW4000. All constructs were cloned in yeast expression vectors with or without a C-terminal tag, as indicated. The respective constructs were transformed into the yeast strain EBY.VW4000 and growth complementation was analyzed on medium containing 2 % glucose or 2 % maltose.

**Figure 4.**
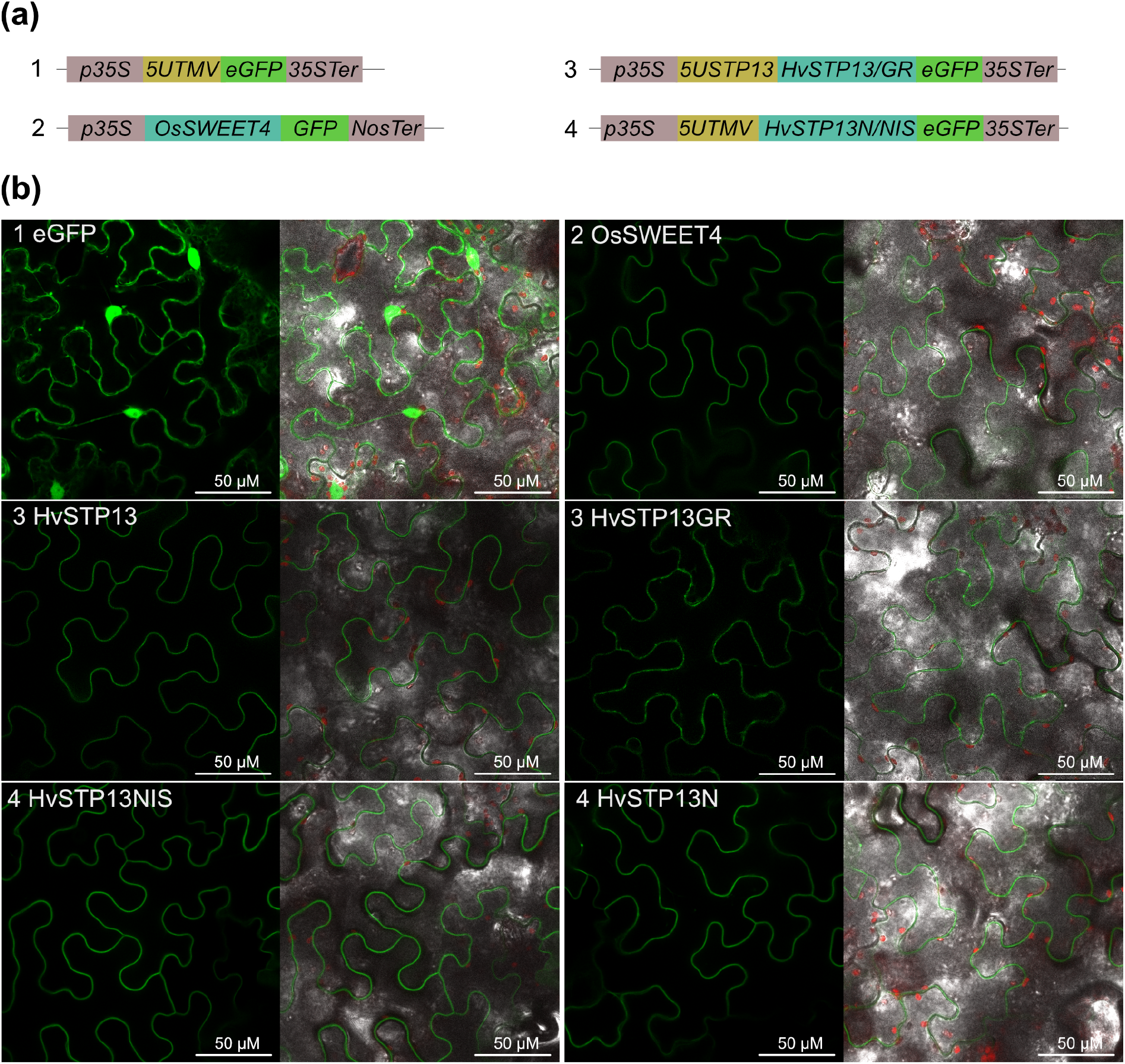
Localization of *HvSTP13* and *HvSTP13* variants *in planta*. **(a)** Overview of the used constructs. **(b)** Constructs were transiently expressed in *N. benthamiana* and visualized 48 hpi by using the Leica SP8 confocal microscope (40 x magnification and 1,5 x zoom). Left panels show eGFP fluorescence and the respective right panels merged pictures (brightfield, chlorophyll autofluorescence and eGFP).

To verify that the lost glucose transport function is not based on a disturbed protein localization, we monitored the subcellular localization of all mutants after transient expression in *N. benthamiana* leaves. The barley HvSTP13 transporter and its HvSTP13GR mutant are postulated to localize to the plasma membrane, but this has not been shown experimentally, so far. Therefore, we cloned all *HvSTP13* variants under control of a short *35S* promoter for transient expression in *N. benthamiana. HvSTP13* and *HvSTP13GR* included the *HvSTP13 5’ UTR*, whereas the *HvSTP13* D41 mutants included the tobacco mosaic virus 5’ UTR that is part of the MoClo module pICH51277 (Figure 4a; (Engler *et al*., 2014)). As positive control we included OsSWEET4, a plasma membrane localized sugar transporter from rice (Sosso *et al*., 2015), while free eGFP served as negative control. Free eGFP localized unspecifically into the nucleus and the cytoplasm, whereas OsSWEET4 exhibited a distinct localization at the edges of each cell that is typical for plasma membrane proteins (Figure 4b). The wildtype HvSTP13 as well as the HvSTP13N and HVSTP13NIS variants displayed a clear localization to the plasma membrane comparable to OsSWEET4. The expression of the HvSTP13GR mutant appeared more patchy, compared to the other transporters, but is also localized predominantly to the plasma membrane (Figure 4b). This demonstrates that all HvSTP13 variants localized to the plasma membrane, suggesting that they did not lose glucose transport activity due to mis-localization or protein degradation. Consequently, the D41 loss of function mutants are promising candidates to analyze their impact for resistance against biotrophic fungal pathogens in the future. As the generation of transgenic or edited plants is time consuming, D41 mutant plants are not available, yet. However, the cytosine base editing approach confirmed that it is at all possible to generate barley plants with a mutation in the genomic sequence of *HvSTP13* for future resistance analysis.

### 2.5 The barley HvSTP13GR mutant confers resistance to a biotrophic fungus

In previous studies the transport-deficient *TaSTP13GR* or *MtSTP13GR* mutants were introduced as a transgene and exerted a dominant negative effect on the wildtype *STP13* copies in the respective genomes, thus conferring resistance in wheat, barley and *Medicago truncatula* (Gupta *et al*., 2021; Milne *et al*., 2019; Moore *et al*., 2015). Presumably, the barley *HvSTP13GR* mutant would also confer resistance to biotrophic fungi when introduced as a transgene. However this has not been analyzed yet.

To analyze the impact of the HvSTP13GR mutant on various pathogen infections, we generated transgenic barley plants that carry the coding sequence of either the *HvSTP13* wildtype, or the *HvSTP13GR* mutant under control of the native *HvSTP13* promoter. To analyze their resistance phenotype, we infected T1 plants of three independent transgenic *HvSTP13* or *HvSTP13GR* lines as well as the wildtype Golden Promise, with the biotrophic rust fungus *Ph* and monitored the infection progress. Two and four days after infection, we harvested leaf sections and stained the fungal structures. At 2 dpi all plants showed an even dispersion of spores on the leaf surface and only minimal differences in their overall developmental stage (Figure S5). Strikingly, at 4 dpi, significant differences in the *Ph* infection development became apparent between *HvSTP13* and *HvSTP13GR* transgenic plants (Figure 5 & 6). Plants from the *HvSTP13* transgenic lines displayed a comparable size of infection sites as the wildtype, with a branched hyphal net, indicating a successful infection (Figure 5). For *HvSTP13GR* transgenic lines, on the other hand, we observed a retarded infection progress, with infection sites being significantly smaller compared to the wildtype or the *HvSTP13* transgenic lines (Figure 5). Moreover, the infection sites never exceeded the formation of only a few haustorial mother cells (Figure 6). Ten days after infection, the leaves of the remaining plants were evaluated regarding their rust pustule formation (Figure 7). For the *HvSTP13*-transgenic and wildtype Golden Promise plants a heavy formation of pustules was observed on the surface of the infected leaves. In contrast, for the *HvSTP13GR*-transgenic plants, only chlorotic spots but no pustules were observed (Figure 7). We confirmed an adequate induction of *HvSTP13* at 48 hpi via qRT-PCR, indicating that the resistance phenotype, observed for the *HvSTP13GR*-transgenic plants, is not based on an impaired expression neither of the genomic nor of the transgenic copy of *HvSTP13* (Figure S6a,b). Instead, it is conceivable that the presence of the transport-deficient HvSTP13GR mutant dominantly affects the function of the wildtype HvSTP13, comparable to the reported resistance mechanism for TaSTP13GR and MtSTP13GR.

**Figure 5.**
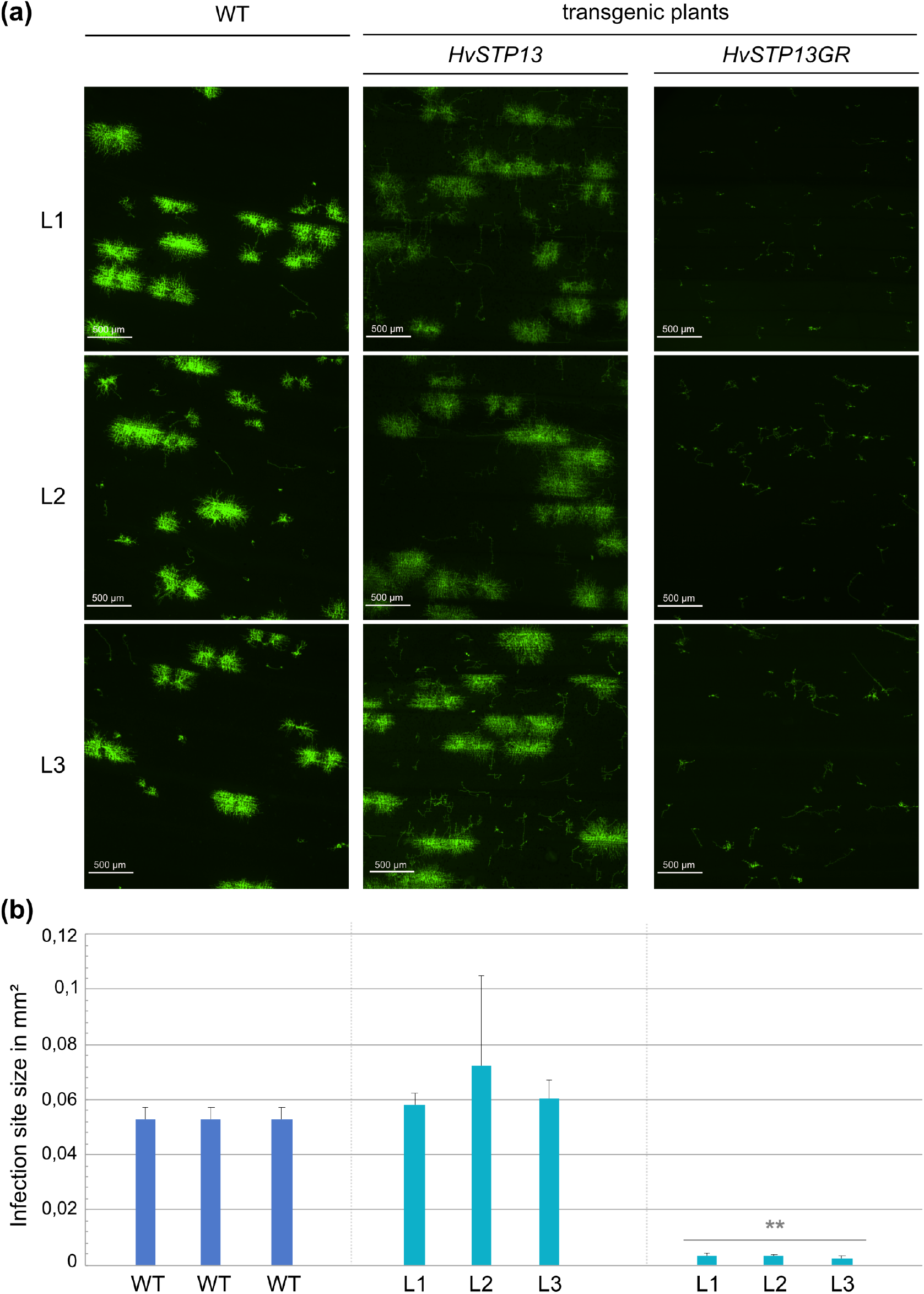
Histological analysis of *Ph* growth in wildtype and transgenic plants at 4 dpi. **(a)** Three wildtype Golden Promise (WT) and three independent *HvSTP13* or *HvSTP13GR* transgenic lines (L1 - L3) were infected with *Ph*. The stained fungal structures were visualized with the Nikon Ti Eclipse at 4 x magnification by using the GFP fluorescence filter. Images are representatives for three independent infection experiments. **(b)** Calculation of infection site size. The values represent the mean of two independent samples and 10 infection sites per sample. The error bars represent the positive standard deviation. Asterisks indicate a significant difference of *HvSTP13GR* transgenic plants from *HvSTP13* transgenic plants and the WT (students t-test; ** p< 0.01).

**Figure 6.**
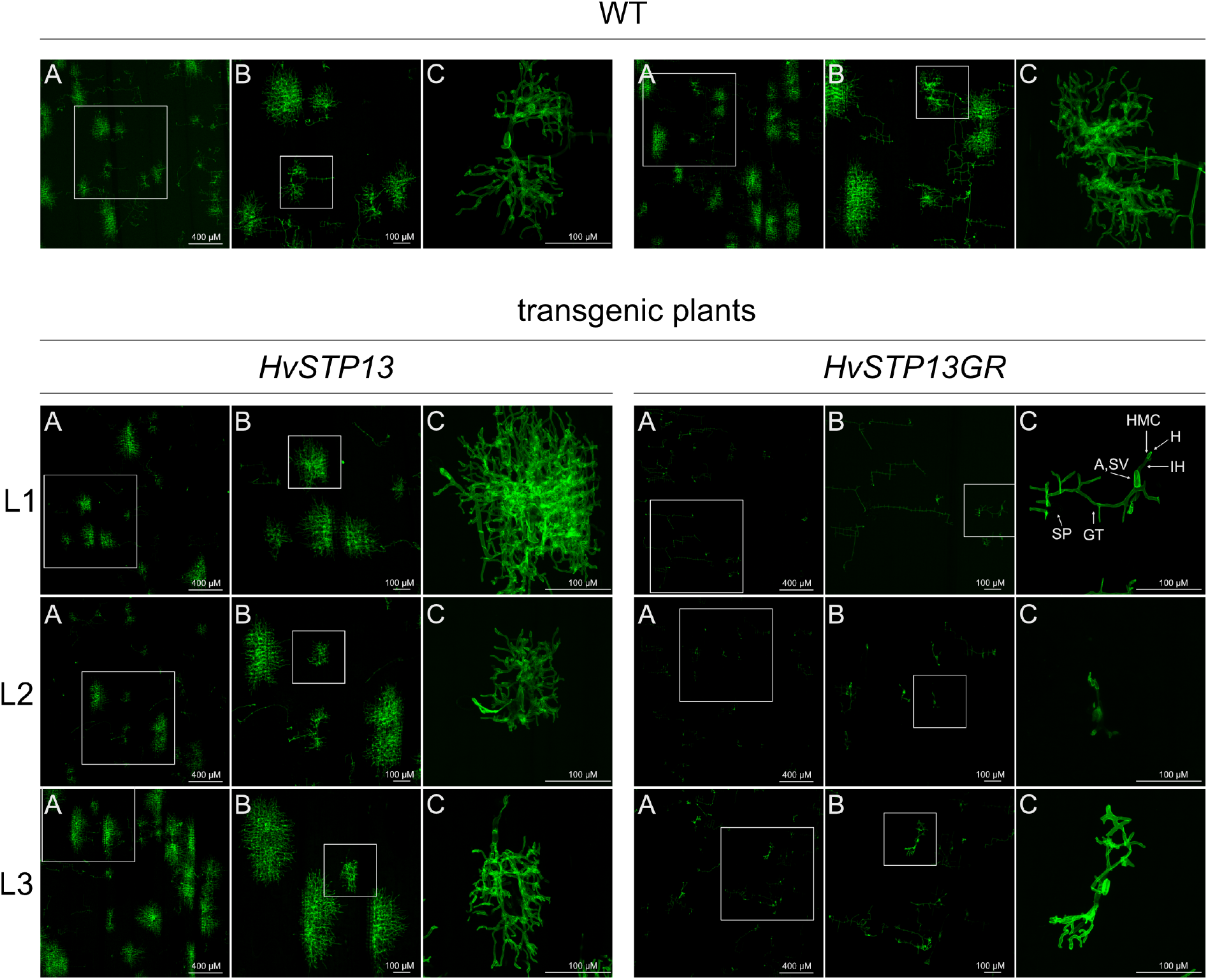
Histological analysis of *Ph* development in wildtype and transgenic plants at 4 dpi. Wildtype Golden Promise (WT) and three independent *HvSTP13* or *HvSTP13GR* transgenic lines (L1 - L3) were infected with *Ph*. The stained fungal structures were visualized with the Leica SP8 confocal microscope at 5 x (A), 10 x (B; Z-stack) and 40 x (C; Z-stack) magnification and GFP fluorescence settings. The respective enlarged area is marked with a grey square. The images are representatives of two biological replicates with at least three independent infection areas each. SP: Spore; GT: Germ tube; A: Appressorium; SV: Substomatal vesicle; IH: Infection hyphe; HMC: Haustorial mother cell; H: Haustorium.

**Figure 7.**
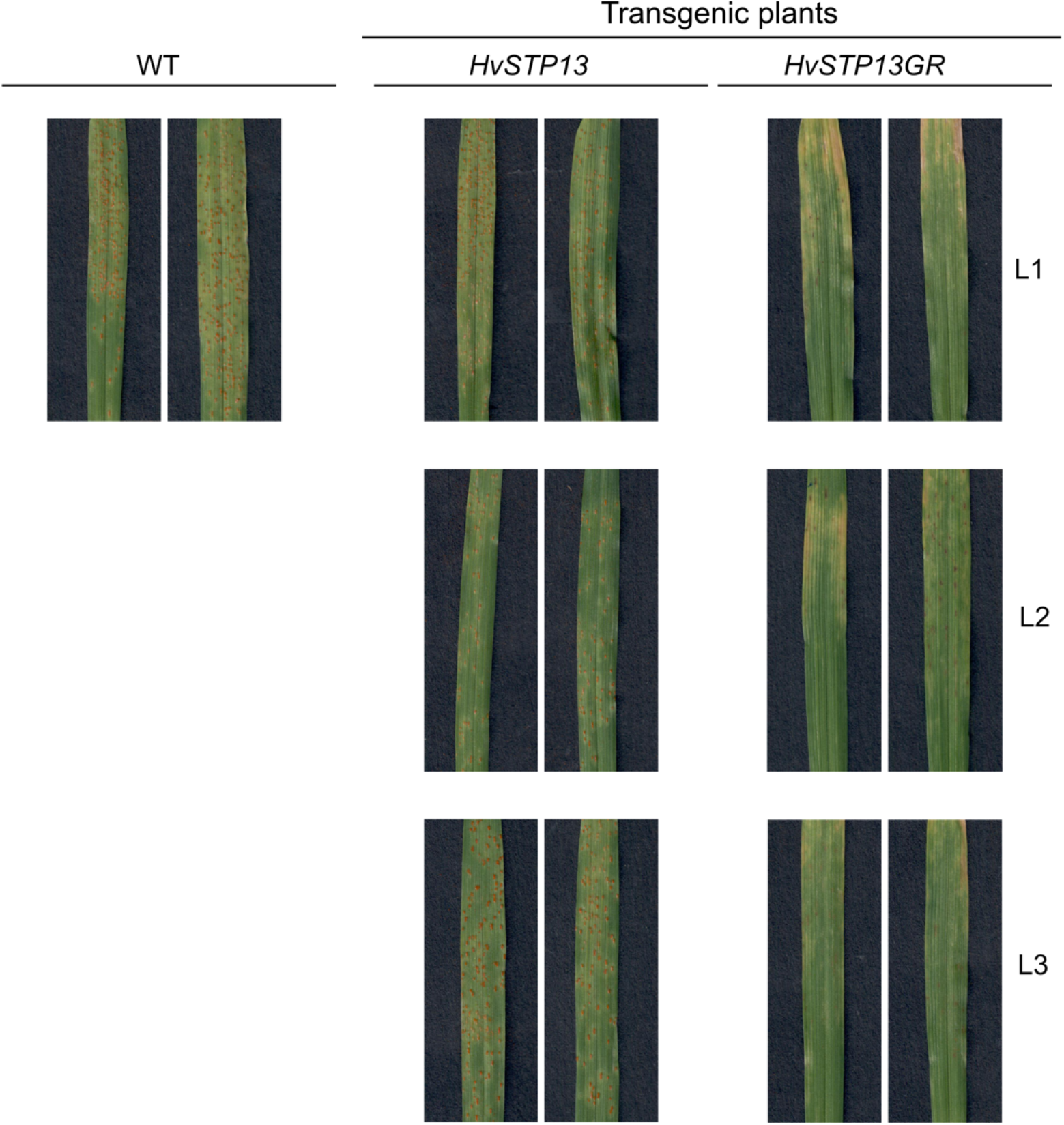
Disease phenotypes of *Ph* infected barley plants at 10 dpi. Wildtype Golden Promise (WT) and three independent *HvSTP13* or *HvSTP13GR* transgenic lines (L1 - L3) were infected with *Puccinia hordei*. The pictures are representative of three independent infection experiments.

## 3 Discussion

In wheat the presence of the glucose transport-deficient TaSTP13GR mutant confers resistance against the two rust fungi *Puccinia triticina* and *Puccinia striiformis* f. sp. *tritici* as well as against the powdery mildew fungus *Blumeria graminis* f. sp. *tritici* (Moore *et al*., 2015). Transferring the TaSTP13GR transgene into barley, mediated resistance against the respective barley rust fungus *Ph* and the powdery mildew fungus *Bgh* (Milne *et al*., 2019). The barley ortholog *HvSTP13* has already been cloned and an equivalent HvSTP13GR mutant has been described as glucose transport-deficient in heterologous yeast studies (Milne *et al*., 2019). However, it has not been analyzed whether the HvSTP13GR mutant confers resistance to biotrophic fungi comparable to TaSTP13GR. Here, we demonstrated that the glucose transport-deficient HvSTP13GR variant confers resistance against the economically relevant rust fungus *Ph*, thus behaving similar to the previously described TaSTP13GR variant (Milne *et al*., 2019). Since this resistance is not based on a gene for gene relation, but possibly on the reduced availability of nutrients for the pathogen, or on increased defense signaling, the manipulation of the *STP13* glucose transporter might present a promising way for the generation of a durable and broad-spectrum resistance - not only for barley but also for other important crops.

Our detailed microscopic analysis of the infection progress for *Ph* demonstrated that the presence of the *HvSTP13GR* transgene does not disturb the initial phase of fungal development until the first haustorial mother cells (HMCs) are formed. However, while the development of *Ph* progresses in *HvSTP13* transgenic and wildtype plants, the fungus revealed an arrested development in the presence of the non-functional HvSTP13GR transporter. Based on our and other findings, different scenarios are conceivable to explain the impact of the glucose transport-deficient STP13GR variant on resistance. On the one hand, the glucose that is imported by STP13 could be an essential carbon source for the fungus, especially in the beginning of the infection. After a spore attaches to the plant surface it germinates and forms infectious structures. In this first developmental phase biotrophic fungi feed from their own nutrient store. But internal stores are limited and the host plant is needed as nutrient source to energize further developmental processes (Divon and Fluhr, 2007). For this, biotrophic fungi form specialized organs called haustoria that invaginate the plant cell membrane and form a tight contact to the plant cytoplasm. There is emerging evidence, that the composition of these organs is unique and provides all components needed for an efficient nutrient or signal transfer between the fungus and the plant (Bozkurt and Kamoun, 2020; Kwaaitaal *et al*., 2017; Polonio *et al*., 2020). Thus, the establishment of this tight interaction platform, to gain nutrients from the host, might be a key step that decides upon failure or success of the disease. Support for this scenario comes from the observation, that diverse fungal pathogens express one or several haustorial glucose transporters that are important for fungal virulence. Furthermore, it was hypothesized that glucose is not only used as energy source but also functions as a signal molecule for fungal growth (Chang *et al*., 2020; van der Linde and Göhre, 2021; Sosso *et al*., 2019; Voegele *et al*., 2001). Fungi secrete invertases, which might further increase the supply with monosaccharides at the interaction surface (Chang *et al*., 2017; Voegele *et al*., 2006). If this glucose supply is hampered by the presence of the transport-deficient STP13GR variant the fungus starves and lacks energy to develop further. Such a disruption of fundamental requirements, would also explain the broad spectrum resistance conferred by the STP13GR mutant against different biotrophic fungi and in different host plants (Gupta *et al*., 2021; Milne *et al*., 2019; Moore *et al*., 2015). On the other hand, various sugars have been shown to regulate cellular processes at multiple levels - including immunity reactions (Bezrutczyk *et al*., 2018). Hence, a blocked glucose import might change the relative glucose concentrations in the apoplast and the cytosol and activate sugar signaling pathways. This shift might be sensed by the intracellular Hexokinase 1 or the extracellular G-protein-coupled receptor RGS1 (regulator of G-protein signaling protein 1). Both proteins are reported to be involved in the activation of defense pathways - including signal transduction, gene expression, stomatal closure, cell wall restructuring or ROS production (Grigston *et al*., 2008; Jing *et al*., 2020; Morkunas and Ratajczak, 2014; Xiao *et al*., 2000; Zhong *et al*., 2019). Here, the inactive STP13GR mutant might result in the activation of those sugar signaling pathways and promote the plants stress response. Furthermore, an altered or gain of function of the TaSTP13GR mutant was discussed by Milne *et al*., 2019 as the basis for the resistance mechanism. Indeed, the G144 to R144 mutation adds an additional positively charged amino acid in close proximity to the proton donor-acceptor pair D42 and R140. This might result in rearrangements of the binding pocket and allow for the binding of an altered substrate that activates an immune response. But, as Huai *et al*., 2020 demonstrated, that also the silencing of the *TaSTP13* transporter increased the resistance against stripe rust, it is still under debate if the loss of the glucose transport activity, a gain of function or a combination of both is the basis for the resistance phenotype of the STP13GR mutants. In future, additional mutants and analysis are needed to elucidate the underlying resistance mechanism.

Sugar transporters play pivotal roles for many physiological processes. Thus, the analysis of sugar transporter mutants is important to decode their impact for sugar partitioning and plant fitness, including pathogen resistance. For the first time, we demonstrated, that cytosine base editing is possible in barley and confirmed this as a suitable approach to generate genomic *HvSTP13* mutants. Our target amino acid D41 is equivalent to D42 in the recently crystalized glucose transporter AtSTP10 (Paulsen *et al*., 2019). In AtSTP10 D42 forms a proton donor-acceptor pair with R142. This mediates the translocation of the H+ proton, an essential prerequisite for the symport of glucose. In HvSTP13 D41 could have a similar function. Our experiments confirm, that the barley HvSTP13 single (N41) and triple (NIS) mutant variants still localized to the plasma membrane but lost their glucose transport activity. This demonstrates the importance of D41 for the sugar transport activity and indicates its possible role in building the proton donor acceptor pair with R140. The D41 mutant is now a promising candidate to dissect whether the transport-deficiency or a new substrate is the cause for the HvSTP13GR mediated resistance. Accordingly, it will be promising to generate D41 mutant plants by using base editing and analyze them in the future.

In wheat, the expression of the closely related ortholog *TaSTP13* is induced by pathogen infection as well as abiotic treatments, including ABA (Huai *et al*., 2020; Moore *et al*., 2015). The comparison of the *HvSTP13* promoter with an equivalent portion of the *TaSTP13* promoter resulted in an identity of 65 % to 68 % (Figure S3b) and it would be interesting in future, to analyze if certain CREs are shared by *TaSTP13* and *HvSTP13*. We demonstrated that the expression of *HvSTP13* is increased upon recognition of bacterial and fungal PAMPs and we identified a promoter region that is probably required for the induced expression. In this promoter region two interesting CREs were predicted, namely two ABREs and a cluster of four W-boxes. ABA, that activates the binding of TFs to ABREs, is mainly related to abiotic stress response, but emerging evidence suggest a cross talk of ABA with pathogen infection. Still, the role of ABA to support or repress immunity reactions largely depends on the pathogen and the phase of infection (Cao *et al*., 2011). Notably, in grapevine, the promoter of the *STP13* ortholog *VvHT5* also employs a unique cluster of ABREs that connects the expression of *VvHT5* to ABA signaling pathways, thus linking it to both, wounding and pathogen infection (Hayes *et al*., 2010). However, whether this observation is connected to PAMP recognition or other effects of pathogen infection is unknown. In comparison, the presence of a W-box cluster in the *HvSTP13* promoter indicates an intriguing link to a PAMP-induced response, as WRKY TFs are key players in downstream processes of the PTI (Chen *et al*., 2019). The interaction of AtSTP13 with FLS2 and its phosphorylation via BAK1 further support such a connection between STP13 activity and PTI signaling, thus rendering WRKYs as promising agents for *HvSTP13* induction (Yamada *et al*., 2016). Increasing research characterized WRKY TFs in important crops as wheat and barley and different roles for WRKY TFs in modulating resistance or susceptibility were presented. For example, HvWRKY10, HvWRKY19 and HvWRKY28 were shown to act as positive regulators in the resistance response of barley to *Bgh* (Meng and Wise, 2012). For wheat it was shown that the transient expression of the barley WRKYs *HvWRKY6, HvWRKY40* and *HvWRKY70* increases the resistance against the biotrophic leaf rust fungus *Puccinia triticina* and a follow-up study confirmed the resistance of wheat expressing HvWRKY6 and HvWRKY70, against *Puccinia striiformis* f. sp. *tritici* and *Blumeria graminis* f. sp. *tritici* (Gao *et al*., 2018; Li *et al*., 2020). Furthermore, HvWRKY23, is activated upon PAMP-recognition by HvCERK1 and probably supports the resistance against the hemi-biotrophic head blight pathogen *Fusarium graminearum* (Karre *et al*., 2017; Karre *et al*., 2019). These examples support the hypothesis that the PAMP-dependent activation of the *HvSTP13* promoter is a result of PTI signaling via WRKY TFs to combat the artificial sink that is generated by pathogens (Pommerrenig *et al*., 2020).

In conclusion, *HvSTP13* might be an important regulatory checkpoint in the multifaceted response of the plant upon both, biotic and abiotic stresses. Our experimental approaches provide novel insights into the activity of HvSTP13 protein variants, their use for breeding, and promoter regions that are relevant for regulating transcription factors. Consequently, our work contributed to further decode the complex regulation network in which STP13 is likely a major player.

## 4 Experimental procedures

### Bacterial infection of barley

The *Pseudomonas syringae* pv. *tomato* strain DC3000 and *Xanthomonas translucens* pv. *hordei* strain UPB820^Rif^ were grown for two days at 28 °C using PSA plates. The strains were resuspended in 10 mM MgCl_2_ adjusted to OD_600_ of 0.2 and inoculated in seven-day-old barley plants. The plants were cultivated in a phytochamber with 16/8 h light and 18/12 °C day/night conditions.

### Fungal infection for *HvSTP13* expression analysis

Fungal infection experiments for *HvSTP13* expression analysis were performed at the Institute for Resistance Research and Stress Tolerance (Federal Research Centre for Cultivated Plants, Julius Kühn Institute (JKI), Quedlinburg, Germany).

### Rust infection

Fresh uredospores for *Puccinia hordei* isolate I-80 were propagated by using the highly susceptible barley cultivar Großklappige as previously described in (Fazlikhani *et al*., 2019). Seven-day-old Golden Promise plants were infected with *Ph* according to (Wehner *et al*., 2019) and cultivated with 16/8h light and 20°C/17°C. *Puccinia striiformis* f. sp. *hordei* isolate 24 was propagated by using Hakai. The infection of Golden Promise plants was performed as described for *Ph* and plants were cultivated with 16/8h light and 16°C/13°C day/night conditions in a greenhouse.

### Powdery mildew infection

Infection was performed as described in (Šurlan-Momirović *et al*., 2016) with the isolate, Freising that was propagated on leaf segments of the susceptible cultivar Igri. Whole plants were infected in the settling tower. Pots with infected plants were placed in a tray with moist sand and covered with transparent plastic boxes. After three days, the air supply was increased by placing small clay pots under the rim of the plastic boxes. Incubation took place at 18°C/15°C 16/8 h day/night cycle.

### Net blotch infection

The mycelium was propagated on PDA plates. One gram of the conidia-bearing mycelium was scraped off the agar plates and 100 ml of sterile water, containing 0.01% Tween 20, was added. The suspension was homogenized for 3 min with a stick blender, filtered through gauze and sprayed evenly onto three-week-old plants. The plants were covered with a plastic hood moistened with water for 48 h. Three days after infection, the plants were exposed to light (16/8 h day/night cycle, 60% humidity) and the temperature was raised from 18 to 20°C according to (Novakazi *et al*., 2019).

### *Ph* infection for resistance analysis in transgenic barley plants

*Ph* infections for resistance analysis were performed at the Department of Plant Biotechnology in Hannover. Fresh uredospores of *Ph* were propagated in climatic chambers (Polyklima, Freysing, Germany) as described before and mixed with activated (10 min, 42°C; 1 h room temperature) frozen spores in a 1:5 ratio. The spores were mixed with clay in a 1:3 ratio. Ten-day-old plants were sprayed with 0.02% Tween 20 using the GANZTON SP180K airbrush system. The spores were applied by using a pipe cleaner. Plastic cups were moistened with 0,02% Tween 20 and used to cover the plants for 48 h. The plants were kept in climatic chambers in darkness for 24 h followed by 16/8 h day/night cycle at 20°C for *Ph*. The infection experiment was repeated three times with 30 plants per transgenic line and 30 wildtype plants, each (120 plants per infection).

### PAMP treatment of barley seedlings

Seven-day-old barley seedlings were inoculated with the PAMPs laminarin (200 µg/µl; Sigma-Aldrich, St. Louis, Missouri, USA), β-1,3-glucan (200 µg/µl; Sigma-Aldrich), chitin (100 µg/ml; Sigma-Aldrich); flg22 (2 µM; Genscript, Piscataway, Township, New Jersey, USA). All PAMPs were dissolved in 10 mM MgCl_2_. 10 mM MgCl_2_ was used as negative control. Plants were cultivated in a phytochamber (16/8 h light; 18/12 °C day/night). Inoculations were done in biological triplicates.

### RNA extraction and gene expression analysis

Total RNA was isolated using the RNeasy Plant Mini Kit (Qiagen, Hilden, Germany). Leaf material was disrupted using the TissueLyser II (Qiagen). On-column DNase digestion was performed using the RNase Free DNase Set (Qiagen). RNA concentration and purity were determined by using the Spark 10M (Tecan, Männedorf, Switzerland). cDNA was synthesized from 2 µg RNA using the Maxima First Strand cDNA Synthesis Kit (Thermo Scientific, Waltham, Massachusetts, USA). All qPCR analyses were performed in technical replicates, including noRT controls, by using the CFX Connect Real-Time PCR Detection System (Bio-Rad, Hercules, California, USA). Reactions contained the Luna Universal qPCR Master Mix (NEB, Ipswich, Massachusetts, USA), 0.25 µM of each primer and 100 ng template cDNA in 20 µl final volume. Primers used for qRT-PCR are listed in Table S3. The ubiquitin-protein ligase, (*HORVU4HrG004090*) served as a reference gene (Hua *et al*., 2015). *HvPR3* (*HORVU1Hr1G052430*; (B., Scheler *et al*., 2016)) served as infection control. Primer efficiencies were determined using a serial cDNA dilution. The fold induction was calculated to the mock control for each time point. Each experiment includes at least two biological replicates and was repeated at least twice.

### Construction of *STP13* modules

The generated level 0 MoClo modules are shown in Figure S7a. The *HvSTP13* (*HORVU1Hr1G052430*) modules were amplified from Golden Promise cDNA or gDNA (Table S3). The yeast *pPMA1* promoter, *ADH2* terminator and *Hxt1* cds were amplified from EBY.VW4000 or already existing constructs (unpublished). Level 1 transcription units were cloned into pICH47742 (plants) or in pAGT572 or pAGT573 (yeast; (U., Scheler *et al*., 2016)). For GUS reporter constructs the MoClo module pICH75111 was used. All multigene constructs for the generation of transgenic plants contained a hygromycine resistance and were generated using the destination vector pAGM8031. For transient expression in *N. benthamiana HvSTP13* variants were cloned under control of the short *35S* promoter. *HvSTP13* and *HvSTP13GR* included the *HvSTP13 5’ UTR* whereas the *HvSTP13* D41 mutants included the tobacco mosaic virus 5’ UTR that is part of the MoClo module pICH51277 (Engler *et al*., 2014).

### Generation of transgenic plants and transgene analysis

Immature embryos of the barley cultivar Golden Promise were transformed with AGL1 according to the protocol of (Harwood *et al*., 2009) with the following modifications. The selection period was reduced to 30 days. 800 mg/L L-cysteine (Hensel *et al*. 2007) and 0.1 mM acetosyringone were added to the co-cultivation medium. After a 10-day transition period, calli were placed on transition medium containing a 50 % reduced hygromycin concentration (25 mg/L) for 4 days. The hygromycin concentration of the regeneration medium was also reduced to 50 %. The transgene status of all regenerated T0 plants and following generations (T1, T2) was analyzed by extracting the gDNA with either the innuPREP Plant DNA Kit (Analytik Jena; Jena, Germany; T0) or the REDExtract-N-AMP plant Kit (Sigma Aldrich; T1, T2). The transgene was detected by amplifying the *Ocs* terminator of the hygromycin resistance cassette (Table S3).

### Analysis of promoter variants via GUS assay

For each GUS reporter construct, three independent T1 transgenic lines with six plants each were used. Two leaf discs (5 mm) were harvested from ten-day-old plants and vacuum infiltrated with 10 mM MgCl_2_, 1 µM flg22 or 100 µg/ml Chitin. The leaf discs were incubated in the phytochamber as described before. After 24 h the leaf discs were frozen in liquid nitrogen and ground by using the TissueLyser II. The GUS activity was measured in the Spark 10M reader as described previously (Kay *et al*., 2007). The measurements were performed in technical replicates. The experiment was performed twice.

### *In planta* localization studies

Four-to six-week-old *N. benthamiana* plants were inoculated with GV3101 strains using a needleless syringe and an OD_600_ of 0.4. The localization of the resulting proteins was analyzed at 2 dpi using the Leica True Confocal Scanner SP8 microscope, equipped with an HC PL APO CS2 40x 1.10 water immersion objective (Leica Microsystems; Wetzlar, Deutschland). Pictures for GFP fluorescence were taken with the HyD detector (488 nm; 502-516 nm) and pictures for chlorophyll autofluorescence with the PMT2 detector (658 – 708 nm). Pictures show a 1.5 x zoom of the 40 x magnification.

### Yeast sugar uptake analysis

All constructs were cloned under control of the *pPMA1* promoter and ADH2 terminator and transformed into EBY.VW4000 (Wieczorke *et al*., 1999). Positively transformed colonies were identified on selective medium with 2 % maltose. Yeast cells were resuspended in sterile H_2_O and adjusted to an OD_600_ of 0.4. Three µl of a serial 10-times dilution were pipetted on selective media with 2 % maltose or 2 % glucose. Plates were incubated at 28 °C for 2 days and growth was documented with the BioRad Chemidoc.

### Histological analysis of fungal growth

Staining of harvested leaves was performed with Alexa Fluor 488 conjugate of WGA according to (Redkar *et al*., 2018) and the stained infection sites were examined using a Nikon Ti Fluorescence microscope (Nikon, Minato, Tokyo Prefecture, Japan). The size of the infection sites was calculated using the object count function of the NIS Elements software. The calculation is based on a one-point threshold definition. With the GFP filter and 10 x magnification, the hue of a characteristic fluorescence signal in the respective infection site was selected to define the threshold for object detection. Based on this calibration, the software calculates the fluorescing area in µm^2^. Detailed pictures were taken with the Leica True Confocal Scanner SP8 microscope using the HyD detector settings for GFP. Z-stacks represent up to 30 panels depending on the growth depth of the fungal structures.

### Cytosine base editing in barley protoplasts

The cytosine base editor was constructed according to Figure S7a,b. The level 0 MoClo modules for the *ZmUbi* promoter, the A3A cytosine base editor, a wheat (*Ta*) codon optimized nCas9 (Ta_nCas9), the uracil glycosylase inhibitor and the Nopaline synthase terminator (pICH41421) were inserted into MoClo level 1 plasmid pICH47742 (Weber *et al*., 2011; Werner *et al*., 2012). The sgRNA expression units were cloned by inserting annealed oligos for the sgRNA spacer sequence into MoClo plasmids (Figure S8). These vectors already contain the *TaU6* promoter and the sgRNA backbone to generate level 0 sgRNA expression plasmids. The sgRNA expression unit was either inserted once or four times into the level 1 MoClo plasmid for position 3 (pICH47751) to generate the 1x sgRNA_D41 or the 4x sgRNA_D41 plasmid. Barley protoplasts were isolated as described in (Shan *et al*., 2014) and transformed with the Ta_nCas9CBE plasmid mixed with one of the two sgRNA plasmids. Each transformation was done twice. 48 hours post PEG-mediated transformation, protoplasts were harvested and gDNA was extracted. The target region was amplified by PCR and six specific nucleotides were added to generate barcoded amplicons for each independent transformation and plasmid combination. The PCR products were pooled and sent for next generation sequencing. The resulting reads were analyzed by using the CRISPR RGEN tool (Hwang *et al*., 2018).

## Acknowledgements

We thank Anna Marthe, Christine Hoppe and Ilona Renneberg for technical assistance and Sebastian Becker and Annekatrin Richter for helpful discussions on the manuscript. We are grateful to Claude Bragard and Ralf Koebnik for providing strain UPB820^R^ and Alain Tissier for providing yeast MoClo plasmids. We thank Dingbo Zhang for cloning the cytosine base editor modules. This work was supported by university core funding only.

## Data availability statement

The data that support the findings of this study are available from the corresponding author upon reasonable request.

## Supporting Information legends

**Figure S1. Transcript level of *HvPR3* after treatment with pathogens and PAMPs**. Barley plants were inoculated with **(a)** *Puccinia hordei* (*Ph*), **(b)** *Puccinia striiformis* f. sp. *hordei* (*Psh*), **(c)** *Blumeria graminis* pv. *hordei* (*Bgh*) and **(d)** *Pyrenophora teres* pv. *teres* (*Ptt*). **(a - d)** *HvPR3* transcript levels were measured via qRT-PCR. Each bar represents three independent biological replicates. Mock = inoculation medium. Error bars represent the standard deviation.

**Figure S2. Transcript level of *HvSTP13* after treatment with pathogens and PAMPs. (a)** Barley plants were inoculated with *Xth* strain UPB820. **(b)** Barley plants were inoculated with the PAMPs laminarin and β-1,3-glucan. **(a) + (b)** *HvSTP13* transcript levels were measured via qRT-PCR. Each bar represents three independent biological replicates. Mock = inoculation medium. Error bars represent the standard deviation.

**Figure S3. Overview of the *HvSTP13* promoter fragments with predicted ABREs, W-boxes and SURE**. The truncated promoter regions, used for generating GUS reporter plants, are marked with purple rectangles and the resulting length is given in nucleotides upstream of the transcriptional start site. The 5’ UTR is marked in orange. Cis-regulatory elements (ABRE, grey; W-box, turquoise; SURE, green) in the respective promoter fragments were predicted using PLACE and PlantCARE. **(b)** A two kb region of the *HvSTP13* promoter was compared with the two kb promoter region of the respective TaSTP13 copys (A, B and D genome). The identity is given in %.

**Figure S4. Cytosine base editing generates HvSTP13 D41 mutants in barley protoplasts. (a)** The sgRNA target sequence, spanning the nucleotides for amino acid D41 (bold), is marked in light green and the PAM in dark green. The plasmid for the cytosine base editor is composed of the A3A cytosine base editor, with a wheat (*Ta*) codon optimized Cas9 (TaCas9CBE). The expression of the sgRNA is controlled by the wheat U6 promoter (*pTaU6*). The sgRNA expression cassette was cloned as single sgRNA construct (1x sgRNA_D41) and as a four times repetition of the sgRNA expression cassette (4x sgRNA_D41). The TaCas9CBE plasmid was mixed with the 1x sgRNA_D41 construct (transformation mix 1) or the 4x_sgRNA_D41 construct (transformation mix 2). The PEG-mediated transformation of barley protoplasts was done twice (1st and 2nd rep). 48 hours post transformation, protoplasts were harvested and gDNA was extracted. The target region was amplified for each transformation independently and specific barcodes were added. The resulting PCR products were pooled and sent for next generation sequencing. **(b) + (c)** The resulting NGS reads were analyzed by using the CRISPR RGEN tool. The total C to T editing efficiency in the target window is given as percentage of the total read number for each PCR product. Sequence variants with an abundance of more than 0,3 % are listed. The resulting amino acid change in comparison to the target sequence is marked in blue.

**Figure S5. Histological analysis of *Ph* growth in wildtype and transgenic plants at 2 dpi**. Wildtype Golden Promise (WT) and three independent *HvSTP13* or *HvSTP13GR* transgenic lines (L1 - L3) were infected with *Ph*. The stained fungal structures were visualized with the Leica SP8 confocal microscope at 5 x (A), 10 x (B; Z-stack) and 40 x (C; Z-stack) magnification by using the GFP fluorescence settings. The respective enlarged area is marked with a grey square. The images are representatives of two biological replicates with at least three independent infection areas each. SP: Spore; GT: Germ tube; A: Appressorium; SV: Substomatal vesicle; IH: Infection hyphe; HMC: Haustorial mother cell; H: Haustorium.

**Figure S6. *HvSTP13* transcript level of WT and transgenic plants 48 hpi with *Ph*. (a) + (b)** The wildtype and three lines of *HvSTP13* or *HvSTP13GR* transgenic plants were inoculated with *Ph. The* transcript levels were measured via qRT-PCR. Each bar represents at least two biological replicates. Mock = inoculation medium. Error bars represent the standard deviation. **(a)** Transcript levels of the genomic *HvSTP13* copy. **(b)** Transcript levels of the *HvSTP13* transgenic copy.

**Figure S7. Generated level 0 MoClo modules and cloning scheme. (a)**. Schematic overview of constructs cloned in this work. All constructs are MoClo compatible. The overhangs created by cloning with BsaI are marked in italics. Pro - promoter; 5’ U – 5’ UTR; CDS1 – coding sequence; CT – C-terminal; 3U+Ter – 3’ UTR + terminator. **(b)** Level 0 modules are combined to a transcriptional unit by cloning into level 1. Level 1 vectors exist for several positions (PX) to generate multigene constructs with BpiI in the next level.

**Figure S8. sgRNA cloning strategy**. sgRNA expression units are cloned by designing and annealing oligos with the 20 nucleotide spacer sequence and flanked by four nucleotides each to generate single stranded overhangs (Step 1). The annealed oligos are inserted into 1, 2, or 4 level 0 empty vectors, respectively (Step 2). These vectors already contain the promoter and the sgRNA backbone (sgBB). The choice of vectors in level 0 determines the sgRNA transcription units that can be combined in level 1. The level 0 vectors can be combined in level 1 with BsaI, to generate one or multiple sgRNA expression units (Step 3). Level 1 empty vectors exist for several positions (PX). The resulting level 1 plasmids can be combined with other level 1 plasmids (e.g. Cas9) to generate multigene constructs in the next level.

**Table S1. PlantCARE prediction results**

**Table S2. PLACE prediction results**

**Table S3. Primer sequences**

