## Supplemental Figures for "The Barley HvSTP13GR mutant triggers resistance against biotrophic fungi"

**Figure S1**

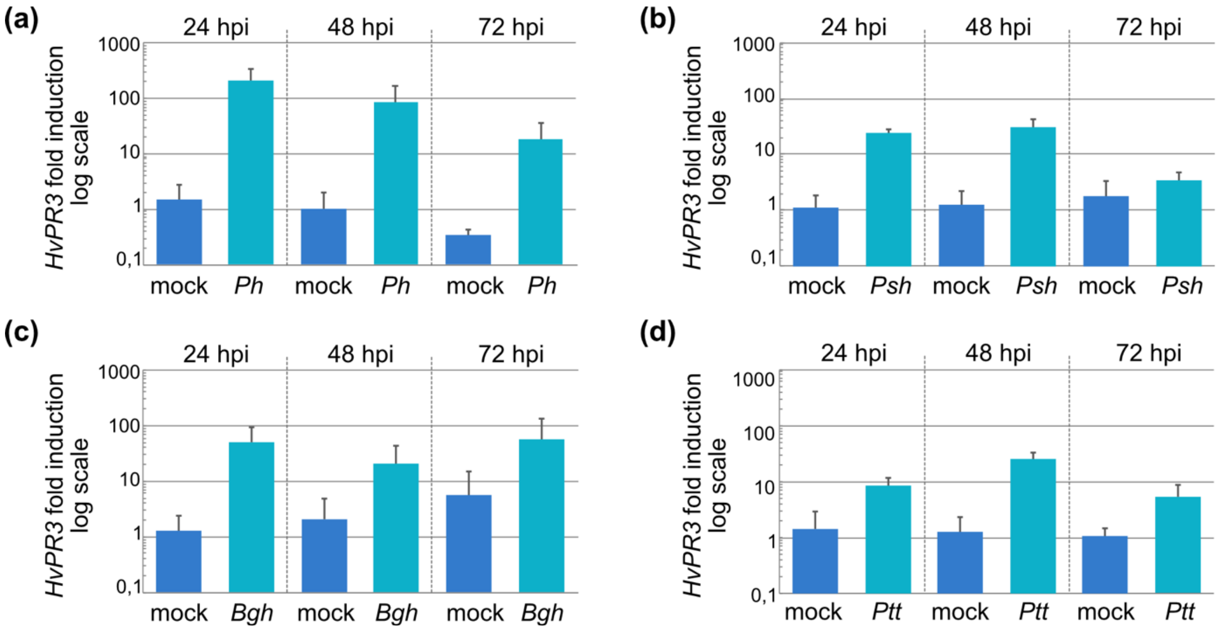


**Figure S1. Transcript level of *HvPR3* after treatment with pathogens and PAMPs.** Barley plants were inoculated with **(a)** *Puccinia hordei* (*Ph*), **(b)** *Puccinia striiformis* f. sp. *hordei* (*Psh*), **(c)** *Blumeria graminis* pv. *hordei* (*Bgh*) and **(d)** *Pyrenophora teres* pv. *teres* (*Ptt*). **(a -d)** *HvPR3* transcript levels were measured via qRT-PCR. Each bar represents three independent biological replicates. Mock = inoculation medium. Error bars represent the standard deviation.

**Figure S2**

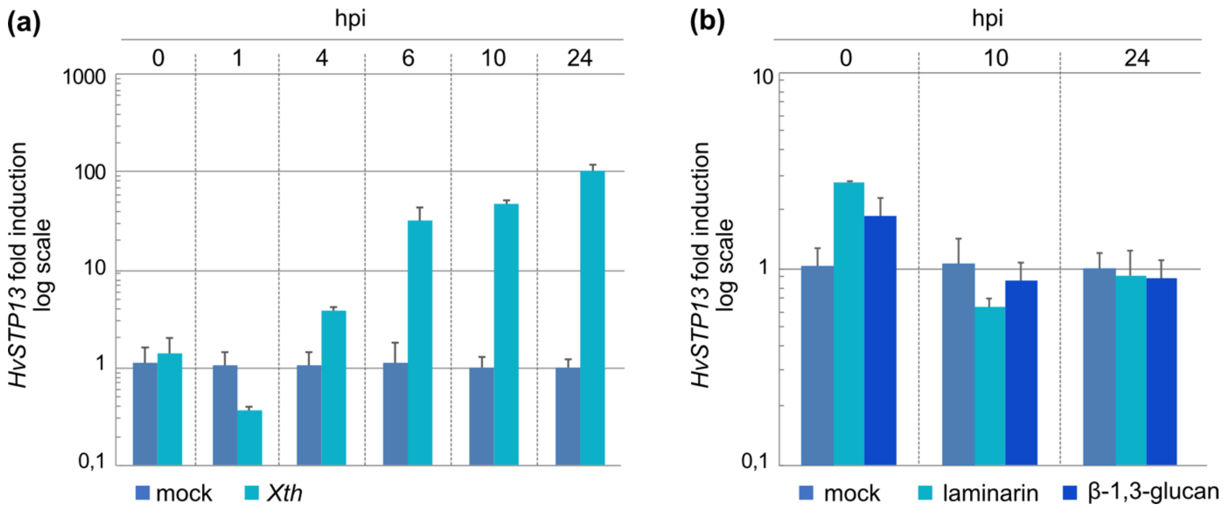


**Figure S2. Transcript level of *HvSTP13* after treatment with pathogens and PAMPs. (a)** Barley plants were inoculated with *Xth* strain UPB820. **(b)** Barley plants were inoculated with the PAMPs laminarin and β-1,3-glucan. **(a) + (b)** *HvSTP13* transcript levels were measured via qRT-PCR. Each bar represents three independent biological replicates. Mock = inoculation medium. Error bars represent the standard deviation.

### Figure S3

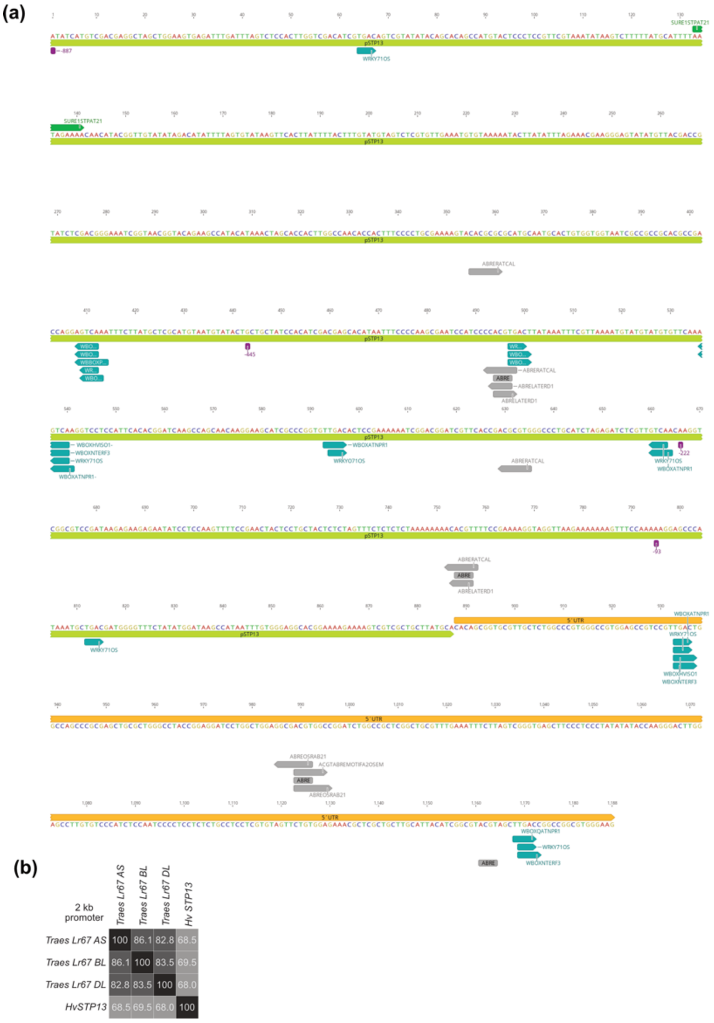


**Figure S3. Overview of the *HvSTP13* promoter fragments with predicted ABREs, W-boxes and SURE. (a)** The truncated promoter regions, used for generating GUS reporter plants, are marked with purple rectangles and the resulting length is given in nucleotides upstream of the transcriptional start site. The 5´UTR is marked in orange. Cis-regulatory elements (ABRE, grey; W-box, turquoise; SURE, green) in the respective promoter fragments were predicted using PLACE and PlantCARE. **(b)** A two kb region of the *HvSTP13* promoter was compared with the two kb promoter region of the respective TaSTP13 copys (A, B and D genome). The identity is given in %.

### Figure S4

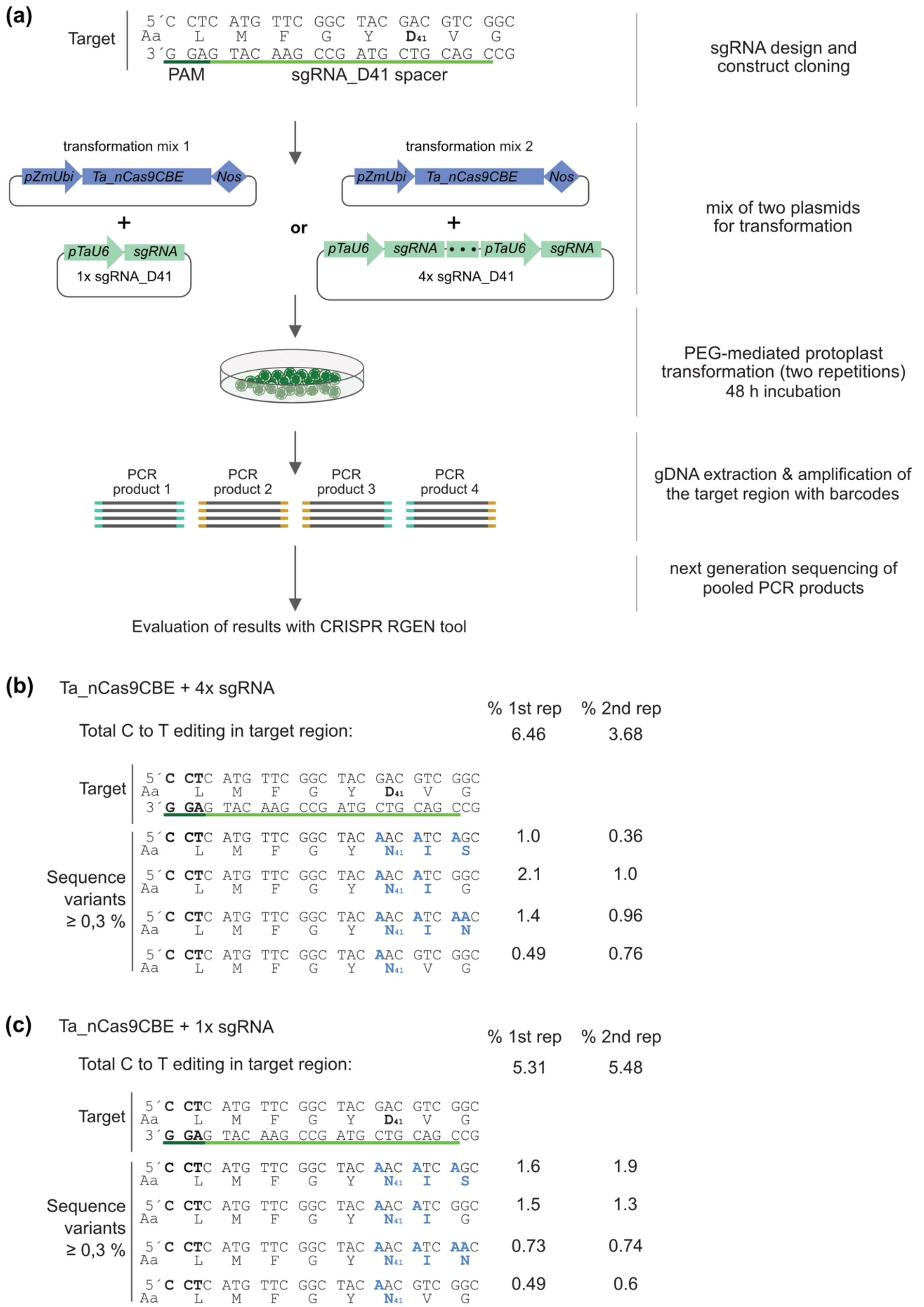


**Figure S4. Cytosine base editing generates HvSTP13 D41 mutants in barley protoplasts. (a)** The sgRNA target sequence, spanning the nucleotides for amino acid D41 (bold), is marked in light green and the PAM in dark green. The plasmid for the cytosine base editor is composed of the A3A cytosine base editor, with a wheat (*Ta*) codon optimized Cas9 (TaCas9CBE). The expression of the sgRNA is controlled by the wheat U6 promoter (*pTaU6*). The sgRNA expression cassette was cloned as single sgRNA construct (1x sgRNA_D41) and as a four times repetition of the sgRNA expression cassette (4x sgRNA_D41). The TaCas9CBE plasmid was mixed with the 1x sgRNA_D41 construct (transformation mix 1) or the 4x_sgRNA_D41 construct (transformation mix 2). The PEG-mediated transformation of barley protoplasts was done twice (1st and 2nd rep). 48 hours post transformation, protoplasts were harvested and gDNA was extracted. The target region was amplified for each transformation independently and specific barcodes were added. The resulting PCR products were pooled and sent for next generation sequencing. **(b) + (c)** The resulting NGS reads were analyzed by using the CRISPR RGEN tool. The total C to T editing efficiency in the target window is given as percentage of the total read number for each PCR product. Sequence variants with an abundance of more than 0,3 % are listed. The resulting amino acid change in comparison to the target sequence is marked in blue.

**Figure S5**

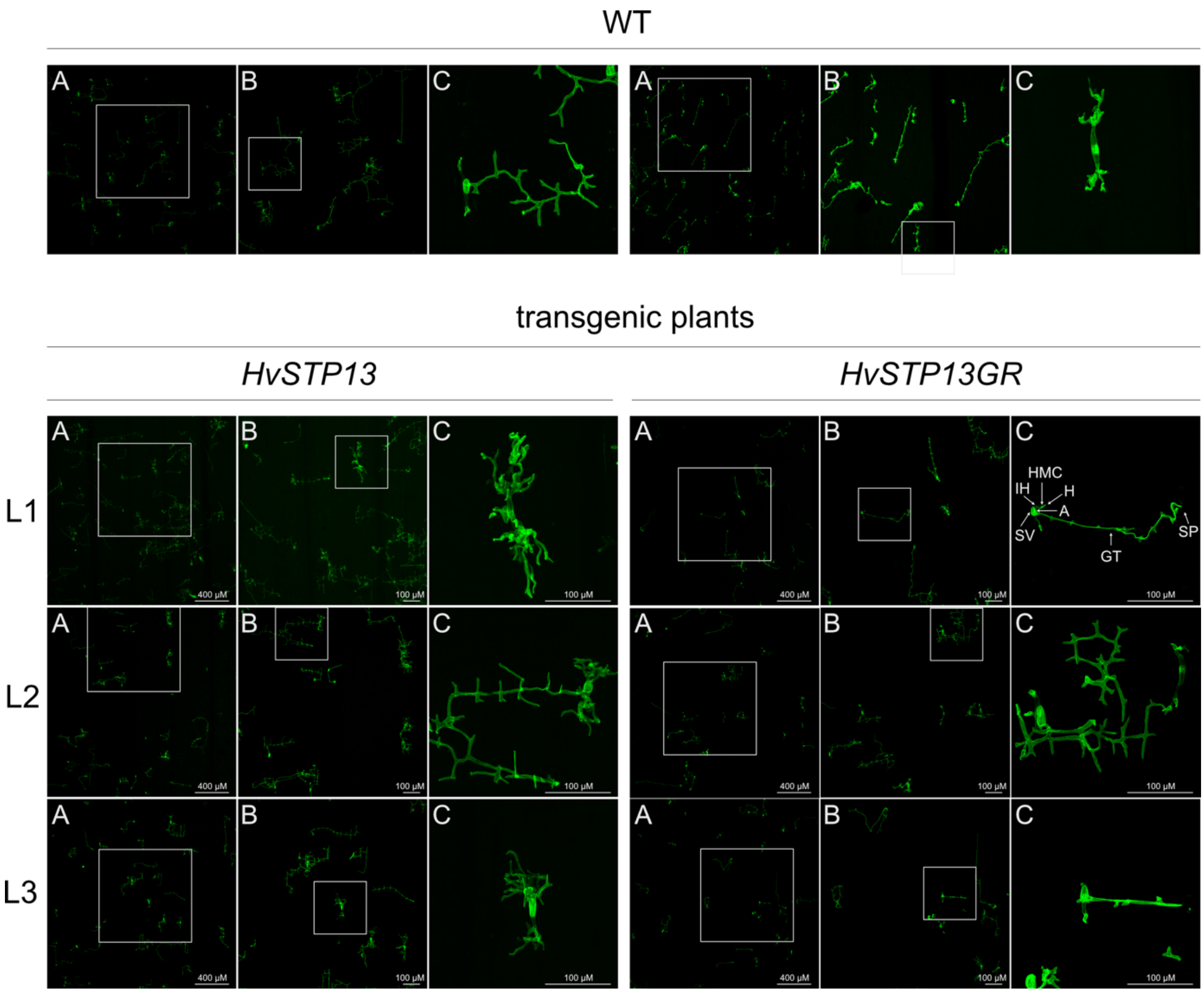


**Figure S5. Histological analysis of *Ph* growth in wildtype and transgenic plants at 2 dpi.** Wildtype Golden Promise (WT) and three independent *HvSTP13* or *HvSTP13GR* transgenic lines (L1 - L3) were infected with *Ph*. The stained fungal structures were visualized with the Leica SP8 confocal microscope at 5 x (A), 10 x (B; Z-stack) and 40 x (C; Z-stack) magnification by using the GFP fluorescence settings. The respective enlarged area is marked with a grey square. The images are representatives of two biological replicates with at least three independent infection areas each. SP: Spore; GT: Germ tube; A: Appressorium; SV: Substomatal vesicle; IH: Infection hyphe; HMC: Haustorial mother cell; H: Haustorium.

**Figure S6**

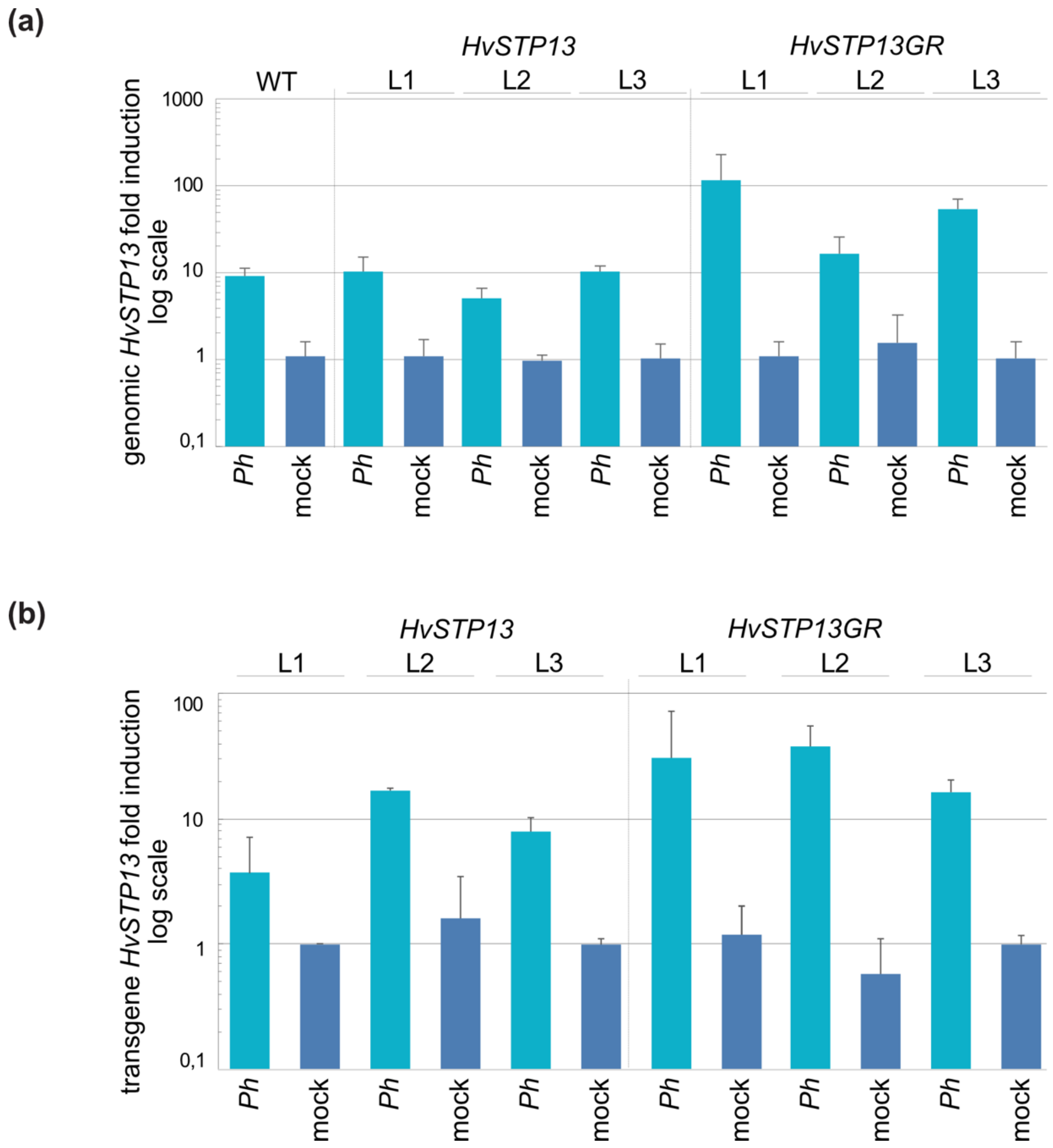


**Figure S6. *HvSTP13* transcript level of WT and transgenic plants 48 hpi with *Ph*. (a) + (b)** The wildtype and three lines of *HvSTP13* or *HvSTP13GR* transgenic plants were inoculated with *Ph*. The transcript levels were measured via qRT-PCR by using specific primers. Each bar represents at least two biological replicates. Mock = inoculation medium. Error bars represent the standard deviation. **(a)** Transcript levels of the genomic *HvSTP13* copy. **(b)** Transcript levels of the *HvSTP13* transgenic copy.

**Figure S7**

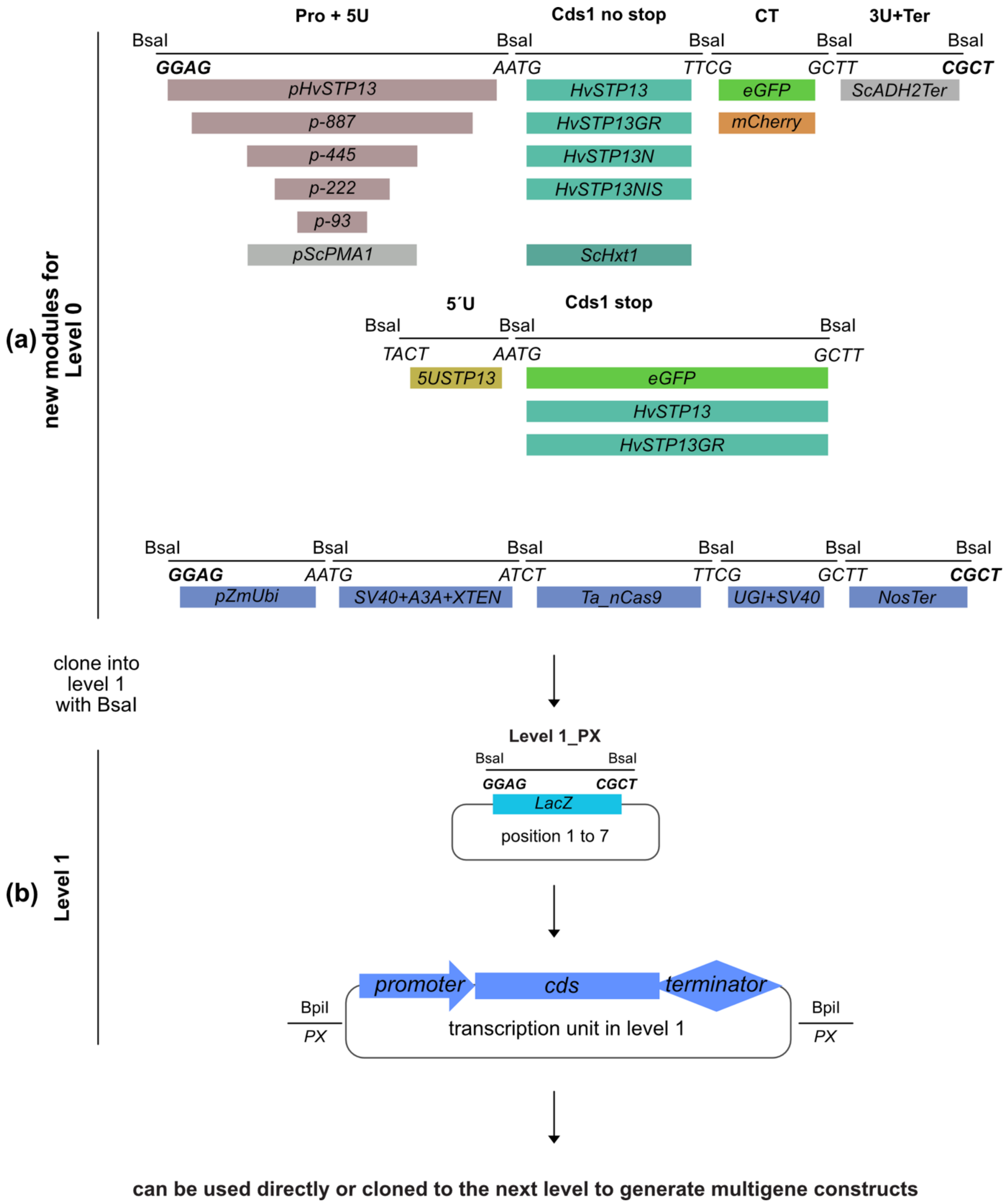


**Figure S7**. **Generated level 0 MoClo modules and cloning scheme. (a)** Schematic overview of the level 0 constructs cloned in this work. All constructs are MoClo compatible. The overhangs created by cloning with BsaI are marked in italics. Pro - promoter; 5´U – 5´UTR; CDS1 – coding sequence; CT – C-terminal; 3U+Ter – 3´UTR + terminator. **(b)** Level 0 modules are combined to a transcriptional unit by cloning into level 1. Level 1 vectors exist for several positions (PX) to generate multigene constructs with BpiI in the next level.

Figure S8

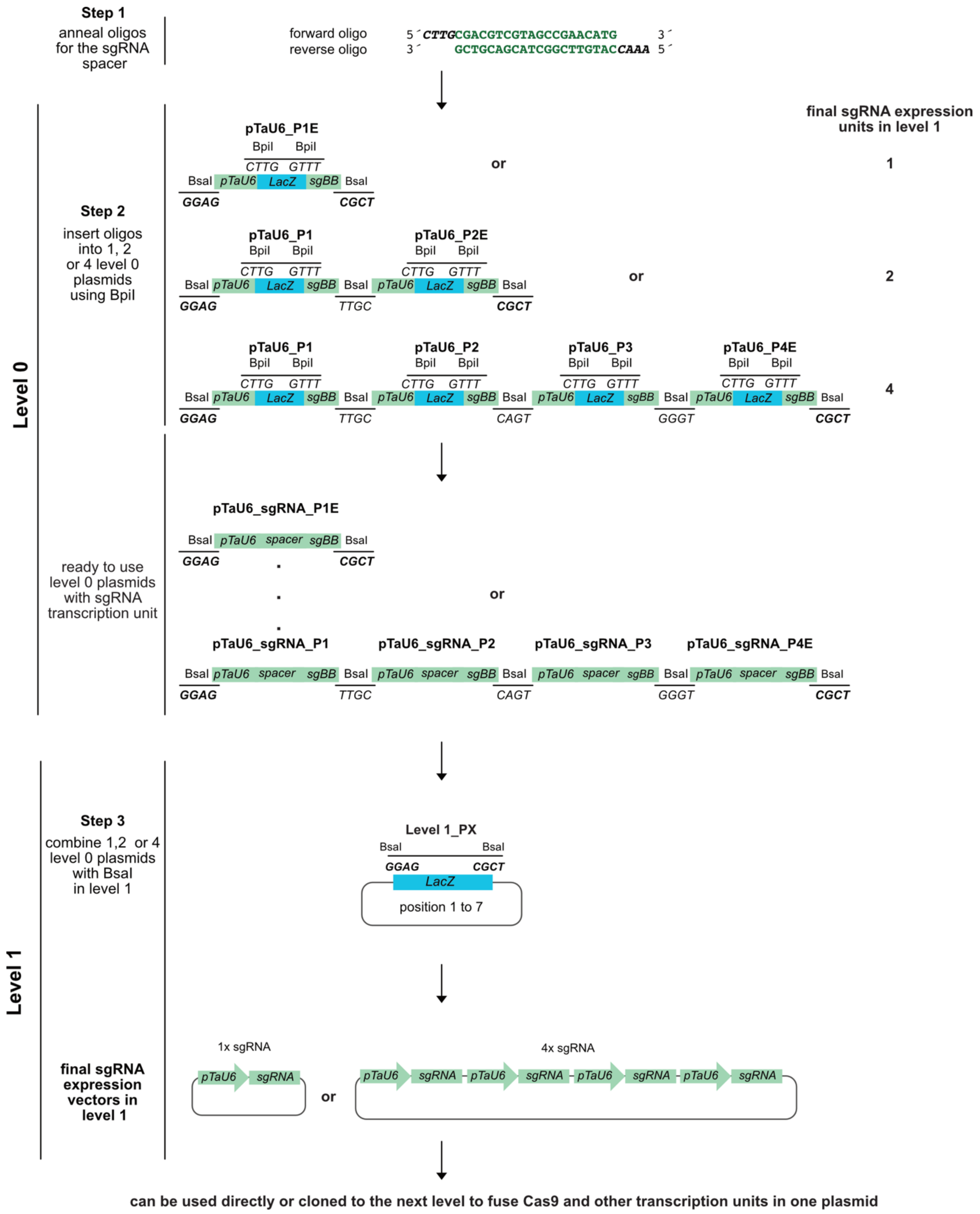


**Figure S8**. **sgRNA cloning strategy.** sgRNA expression units are cloned by designing and annealing oligos with the 20 nucleotide spacer sequence and flanked by four nucleotides each to generate single stranded overhangs (Step 1). The annealed oligos are inserted into 1, 2, or 4 level 0 empty vectors, respectively (Step 2). These vectors already contain the promoter and the sgRNA backbone (sgBB). The choice of vectors in level 0 determines the sgRNA transcription units that can be combined in level 1. The level 0 vectors can be combined in level

1 with BsaI, to generate one or multiple sgRNA expression units (Step 3). Level 1 empty vectors exist for several positions (PX). The resulting level 1 plasmids can be combined with other level 1 plasmids (e.g. Cas9) to generate multigene constructs in the next level.
