## Supplementary material for "The Barley HvSTP13GR mutant triggers resistance against biotrophic fungi": Table S3

**Table S3 Primer Sequences**

| **Name** | **Sequence^a^ 5´- 3´** | **Used for** |
| --- | --- | --- |
| HvSTP13_cds1_F | TTT**GAAGAC**TT**AATG**CCGGGCGGAGGGTTC | Cloning |
| HvSTP13_4NoSTOP_R | TT**GAAGAC**AA**CGAA**CACGGTGGCGTTCTTGCC | Cloning |
| HvSTP13_4Stop_R | TT**GAAGAC**AA**AAGC**TTAGACGGTGGCGTTCTTGC | Cloning |
| HvSTP13_Pro5U_F | TT**GAAGAC**TT**GGAG**CGAATCAATCTTTTCTGTTTATTGATCTG | Cloning |
| HvSTP13_Pro5U_R | TT**GAAGAC**AA**CATT**CTTCCCACGCCGGCCGG | Cloning |
| pSTP13_-93_F | TT**GGTCTC**T**GGAG**AAGGAGCCCATAAATGCTGACG | Cloning |
| pSTP13_-222_F | TT**GGTCTC**T**GGAG**AAGGTCGGCGTCCGATAAGAG | Cloning |
| pSTP13_-445_F | TT**GGTCTC**T**GGAG**GCTGCTATCCACATCGACGAG | Cloning |
| pSTP13_-887_F | TT**GGTCTC**T**GGAG**ATATCATGTCGACGAGGCTAGCTG | Cloning |
| pSTP13_Pro5U_Bsa_R | TTT**GGTCTC**A**CATT**CTTCCCACGCCGGCC | Cloning |
| STP13_5U_F | TT**GGTCTC**T**TACT**CACAGCGGTGCGTTGCTC | Cloning |
| STP13_5U_R | TT**GGTCTC**A**CATT**CTTCCCACGCCGGCC | Cloning |
| pPMA1_Pro5U_R | TT**GGTCTC**A**CATT**ATTGATATTGTTTGATAATTAAATC | Cloning |
| pPMA1_Pro_F | TT**GGTCTC**T**GGAG**CTCAGCTTTGCTAAAGTGCAAAAAG | Cloning |
| ADH2Ter_F | TT**GGTCTC**T**GCTT**AAGCTTTGGACTTCTTCGCCAG | Cloning |
| ADH2Ter_R | TT**GGTCTC**A**AGCG**GGCCGGTAGAGGTGTGG | Cloning |
| STP13_42N_F | AACGTCGGCATCTCAGGCGG | STP13 mutagenesis |
| STP13_42NIS_allR | GTAGCCGAACATGAGGCCG | STP13 mutagenesis |
| STP13_42NIS_F | AACATCAGCATCTCAGGCGGGGTAAC | STP13 mutagenesis |
| HvSTP13_GR_F | AGGTGTGGCGTCGGCTTCGC | STP13 mutagenesis |
| HvSTP13_GR_R | GAGCAGGATCCTGCCGACG | STP13 mutagenesis |
| HvSTP13_qRT_R | TCTTGGCATGTCAGCAGTGT | qRT genomic copy (Milne *et al*., 2019) |
| HvSTP13_qRT_F | CTCCGTCCTACTCCTATGTAA | qRT genomic copy (Milne *et al*., 2019) |
| HvSTP13_qRT_TransF | GGGTGCTCGTCATGTCCGTG | qRT transgene |
| HvSTP13_qRT_TransR | CATCGCAAGACCGGCAACAG | qRT transgene |
| UPL_qRT_F | CTGAAGAGTTAGGCGGGAAA | qRT (Hua *et al*., 2015) |
| UPL_qRT_R | TCGCATGAACGTAGTGCAA | qRT (Hua *et al*., 2015) |
| PR3_qRT | CCAACACCTTTCCGGGCTTCG | qRT (Scheler *et al*., 2016) |
| PR3_qRT | CAATTGGATGGGTCCCCGTCC | qRT (Scheler *et al*., 2016) |
| OcsTer_F | GAGATATGCGAGAAGCCTATGATCG | Transgene detection |
| OcsTer_R | GACGGCCAATACTCAACTTCAAGG | Transgene detection |
| Gus 5´RACE/SK | GAACTGATCGTTAAAACTGCCTG | Transgene detection^b^ |
| AS_STP13D42_FI1 | GAG AAC AGT TCG AGG CCA AGA TCA CG | Amplicon sequencing |
| AS_STP13D42_RI1 | ACA GGC GCT GCA ACG GAA GTA TAC GC | Amplicon sequencing |
| As_STP13D42_I3F | TCTCCTAGTTCGAGGCCAAGATCACG | Amplicon sequencing |
| As_STP13D42_I3R | CGAATGGCTGCAACGGAAGTATACGC | Amplicon sequencing |

**^a^** BpiI or BsaI recognition sites and resulting overhangs are marked in bold

**^b^** Primer was used in combination with the forward primer of the respective fragment
